# An Entropic Safety Catch Controls Hepatitis C Virus Entry and Antibody Resistance

**DOI:** 10.1101/2020.11.11.377218

**Authors:** Lenka Stejskal, Mphatso D. Kalemera, Machaela Palor, Lucas Walker, Tina Daviter, William D. Lees, David S. Moss, Myrto Kremyda-Vlachou, Zisis Kozlakidis, William Rosenberg, Christopher J. R. Illingworth, Adrian J. Shepherd, Joe Grove

**Affiliations:** Institute of Immunity and Transplantation, Division of Infection and Immunity, The Royal Free Hospital, University College London, UK; Institute of Structural and Molecular Biology, Birkbeck College, University of London, London, UK; Shared Research Facilities, The Institute of Cancer Research, London, UK; Division of Infection and Immunity, University College London, UK; International Agency for Research on Cancer, World Health Organization, Lyon, France; Division of Medicine, Institute for Liver and Digestive Health, University College London, UK; MRC Biostatistics Unit, University of Cambridge, UK; Department of Genetics, University of Cambridge, UK; Institut für Biologische Physik, Universität zu Köln, Germany

**Keywords:** virus entry, protein disorder, hepatitis C, hypervariable region-1

## Abstract

E1 and E2 (E1E2), the entry proteins of Hepatitis C Virus (HCV), are unlike that of any other virus yet described, and the detailed molecular mechanisms of HCV entry/fusion remain unknown. Hypervariable region-1 (HVR-1) of E2 is a putative intrinsically disordered protein tail. Here, we demonstrate that HVR-1 has an autoinhibitory function that suppresses the activity of E1E2 on free virions; this is dependent on its conformational entropy. Crucially, to allow entry, this mechanism is turned off by host receptor interactions at the cell surface. Thus, HVR-1 is akin to a safety catch on E1E2 activity. Mutations that reduce conformational entropy in HVR-1, or genetic deletion of HVR-1, turn off the safety catch to generate hyper-reactive HCV that exhibits enhanced virus entry but is thermally unstable and acutely sensitive to neutralising antibodies. Therefore, the HVR-1 safety catch controls the efficiency of virus entry and maintains resistance to neutralising antibodies.

## Introduction

The entry of all enveloped viruses is dependent on their fusion machinery. These typically come in the form of glycoprotein complexes on the virion surface which act to bring host and viral membranes together (Grove and Marsh, 2011; Kielian and Rey, 2006). Being prominently displayed, these proteins are also the principal target of neutralizing antibodies (nAbs). Antibody binding often results in the inactivation of entry machinery, either through blocking receptor interactions, preventing conformational rearrangements or destabilisation (McCoy, 2018; Munro et al., 2014; Walls et al., 2019). Investigating the various mechanisms by which viruses achieve entry has uncovered novel aspects of fundamental cell biology and guided the rational design of therapies and vaccines (Burton et al., 2012; Crank et al., 2019; Fédry et al., 2017; McLellan et al., 2013).

Convergent evolution has led diverse viruses to develop strikingly similar entry apparatus, which can be broadly categorized as class-I, II or III fusion machines. For example, HIV, Ebolavirus, Influenza and Coronaviruses all exhibit structurally similar class-I fusion machinery (Rey and Lok, 2018). However, based on current structural information, the entry proteins of Hepatitis C Virus (HCV), E1 and E2 (E1E2), do not possess the hallmarks of any previously described fusion proteins (El Omari et al., 2014; Flyak et al., 2018; Khan et al., 2014, 2015; Kong et al., 2013; Tzarum et al., 2019); therefore, E1E2 may represent the prototype of a new class of fusion machinery and is likely to exhibit completely novel features. Moreover, a complete structure-to-function understanding of E1E2 will expedite ongoing HCV vaccine development.

Given E1E2 are unlike any known viral entry machine, the molecular mechanics of HCV entry will have to be discovered de novo. Nonetheless, there are general concepts that can be applied to all viral fusion proteins. The function of viral fusion machinery is to tightly juxtapose viral and host membranes, allowing them to fuse. Bringing two membranes together requires an energy input to expel any intervening molecules, such as the hydration shell around the lipid bilayers. Fusion proteins provide this energy input by undergoing dramatic refolding during virus entry. However, this conformational transition requires careful regulation; the process is irreversible, therefore, premature triggering (prior to docking at a cellular membrane) will inactivate the machinery; conversely, failure to trigger during virus entry will prevent fusion. Consequently, viral fusion proteins rely on specific molecular cues, provided by the host, to ensure timely and effective activation. Therefore, viral entry proteins can be viewed as being spring-loaded conformational machines with molecular checkpoints to prevent premature triggering (Kielian and Rey, 2006; Rey and Lok, 2018).

HCV entry involves at least four host factors: CD81, scavenger receptor B-1 (SR-B1), claudin-1 (CLDN1) and occludin (OCLN). Additionally, epidermal growth factor receptor signalling contributes to receptor-complex formation and particle endocytosis, after which pH-dependent fusion occurs in the early endosome (Baktash et al., 2018; Evans et al., 2007; Lupberger et al., 2011; Pileri et al., 1998; Ploss et al., 2009; Scarselli et al., 2002). It is thought that only two host factors, SR-B1 and CD81, interact directly with HCV, via the major glycoprotein E2 and current evidence suggests that the minor glycoprotein, E1, contains the fusogen (Hu et al., 2020; Perin et al., 2016). Whilst there is a good structural understanding of the E2 ectodomain and partial characterisation of E1, how they assemble and function together is poorly understood (Cao et al., 2019). In particular, the molecular consequences of E1E2 interaction(s) with receptors and how this relates to the stepwise priming and triggering of the HCV fusion mechanism remains unknown.

Here, we demonstrate that genetic substitutions in E2 can switch the E1E2 entry machinery into a hyper-reactive state, this increases particle infectivity by enhancing the efficiency and kinetics of virus entry. However, hyper-reactive HCV is thermally unstable and acutely sensitive to nAbs suggesting a high propensity for inactivation of E1E2, presumably through the premature triggering of fusion activity. These mutations also confer a low dependency on SR-B1, indicating a role for this receptor in regulating the reactivity of E1E2. We recently demonstrated that hypervariable region-1 (HVR-1), which contains the SR-B1 binding site, is a putative intrinsically disordered protein tail that displays high conformational entropy (Stejskal et al., 2020). Molecular dynamic simulations and biophysical analysis reveal that hyper-reactive mutants exhibit stabilisation of HVR-1. Based on these data we hypothesised that the structural disorder of HVR-1 exerts an autoinhibitory effect on E1E2, and that the role of SR-B1 is to bind and stabilise HVR-1, removing autoinhibition. Therefore, HVR-1 acts much like a safety catch on a firearm, controlling the propensity of E1E2 to trigger. Turning off the safety catch at the cell surface is the first stage of HCV entry. Consistent with this, genetic deletion of the HVR-1 safety catch, switches HCV into a hyper-reactive state. Moreover, we demonstrate that antibody selection ensures that the safety catch remains engaged, therefore maintaining low E1E2 reactivity and high resistance to antibody-mediated neutralisation.

## Results

### HCV explores evolutionary pathways to optimise virus entry

We established a continuous culture HCV (J6/JFH HCVcc) in Huh-7.5 cells and monitored viral evolution by next-generation sequencing (NGS). By day 42 we detected various substitutions throughout the genome (prior to this time, no mutations reached our 5% frequency threshold). The viral entry glycoproteins, E1E2, were particularly enriched for non-synonymous substitutions when compared to other coding regions (Figure S1A), suggesting that virus entry may be undergoing adaptive optimisation. These substitutions were largely located towards the C-terminus of E1 and the N-terminus of E2, the latter being important for receptor and nAb interactions (Figure S1B-D) (Tzarum et al., 2018). Some of the substitutions occurred at highly-conserved sites (e.g. V371A, G406S, S449P), suggesting that in vitro replication does not recreate the evolutionary constraints that are usually found in vivo; we note that a critical difference between these settings is the complete absence of an adaptive immune response in vitro. From day 42 onward we observed the emergence of a mutant lineage with sequential fixation of I438V and A524T (found in the front layer and CD81 binding loop of E2, respectively, Figure S1E), and concomitant loss of other variants from the population (e.g. I262L, Figure S2A). Introduction of these sequential mutations, by reverse genetics, resulted in a stepwise increase in HCV infectivity (Figure 1A), suggesting enhanced entry efficiency. We evaluated the entry pathway of these mutants by infecting cells in which the critical HCV entry factors (CD81, SR-B1, CLDN1 and OCLN) had been knocked out by CRISPR/Cas9 gene editing (Figure 1B). WT and mutant viruses were equally dependent on CD81, CLDN1 and OCLN, whereas the requirement for SR-B1 decreased in the mutant viruses in a manner that mirrored viral titre, as evidenced by infection of SR-B1 KO cells. This demonstrates that the efficiency of HCV entry is tightly linked to SR-B1 dependency.

**Fig. 1.**
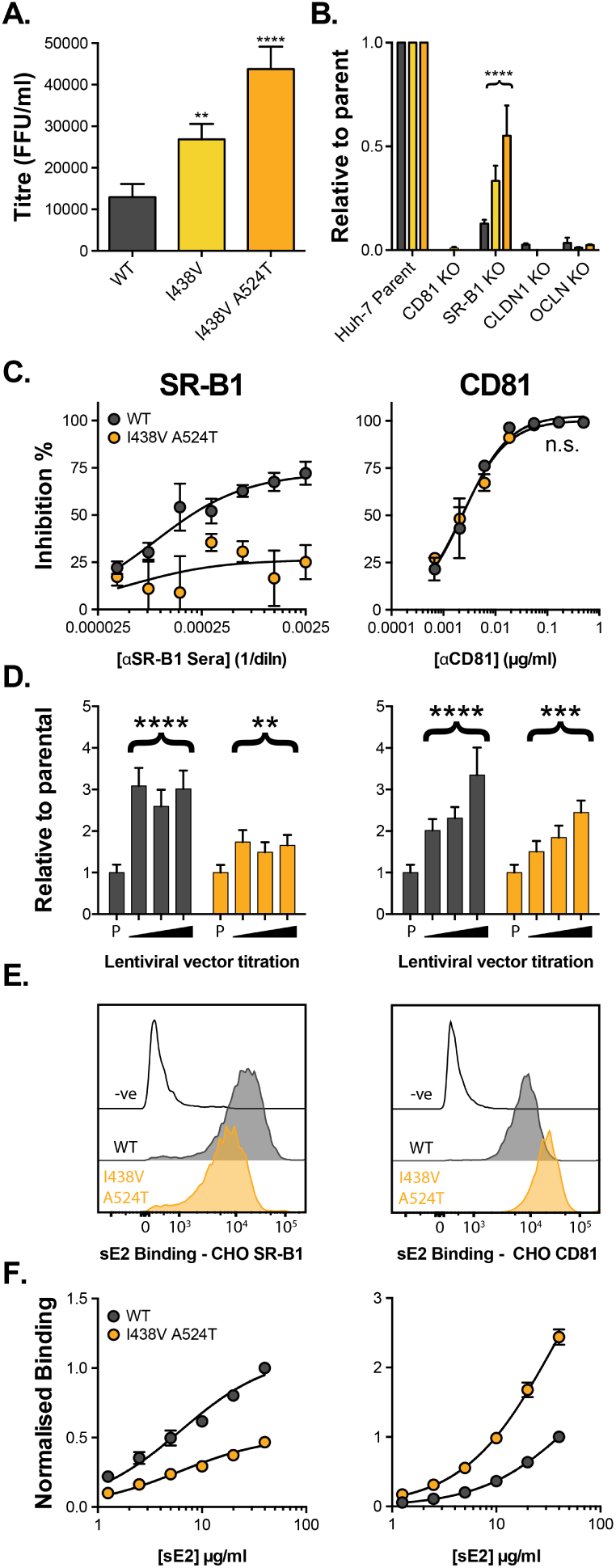
HCV evolves to optimise entry through altered receptor dependency. J6/JFH HCVcc mutants were isolated following continuous propagation in Huh-7.5 cells. **A**. Infectivity of WT, I438V and I438V A524T HCVcc, expressed as foci forming units per ml. Values were normalised for input particle numbers. **B**. WT and mutant HCV infection of parental Huh-7 cells and those CRISPR/Cas9 engineered to knock out the stated HCV entry factor. To aid direct comparison of each mutant, infection values have been expressed relative to that observed in parental Huh-7 cells. Values represent the mean of three independent experiments **C**. Huh-7.5 cells were treated with anti-SR-B1 serum (left) or anti-CD81 mAb (right) to limit receptor availability. Infection by WT and I438V A524T HCVcc is expressed as % inhibition relative to infection of untreated cells. Data points represent the mean of three independent experiments. **D**. To increase receptor availability, Huh-7.5 cells were transduced with a serial dilution of lentiviral vectors encoding either SR-B1 (left) or CD81 (right). Infection by WT and I438V A524T HCVcc is expressed relative to infection of parental cells (P). Example data from one representative transduction is shown. **E**. & **F**. CHO cells were transduced to express exogenous human SR-B1 (left) or CD81 (right), to which WT and mutant sE2 binding was assessed by flow cytometry. Upper plots (E.) provide representative cytometry histograms of sE2 binding to transduced or parental (-ve) CHO cells. Lower plots (F.) display quantification of sE2 binding, with values normalised to WT binding at 40*µ*g/ml. Data points represent the mean of three independent experiments. In all cases, error bars indicate standard error of the mean, asterisks denote statistical significance (ANOVA, GraphPad Prism). Curve fitting performed with a hyperbola function and the curves compared to confirm significance difference (F-test, GraphPad Prism); ns denotes the lack of significant difference.

We also observed the acquisition of an additional mutation in this lineage (M356V, in E1), which conferred a further increase in infectivity (Figure S2B), but no further change in receptor dependency (Figure S2D). Also, viruses bearing either A524T or M356V single mutations were phenotypically identical to WT (Figure S2C & D), indicating a hierarchical interdependence between the mutations. Thus, the double E2 mutations (I438V A524T) were necessary and sufficient to confer altered receptor dependency, therefore we focussed our investigation on this virus.

Thus far, only SR-B1 and CD81 have been proven to interact directly with the HCV entry machinery, via E2 (Pileri et al., 1998; Scarselli et al., 2002). The precise molecular basis of E2-receptor interactions have yet to be defined at the structural level; nonetheless, mutational and antibody blocking experiments have demonstrated that SR-B1 binding occurs via the N-terminal HVR-1, whilst the CD81 binding site is thought to be composed of three discontinuous regions (antigenic site 412 [AS412], the front layer, and the CD81 binding loop) (Kong et al., 2013; Owsianka et al., 2006; Scarselli et al., 2002). Previous work, including our own, suggests that SR-B1 is the initial receptor for HCV and is likely to prime subsequent stages of entry (including interaction with CD81) via an unknown mechanism (Augestad et al., 2020; Evans et al., 2007; Kalemera et al., 2019).

We evaluated the relationship of WT and I438V A524T HCV with SR-B1 and CD81 using a variety of techniques. First, we performed receptor blockade using antibodies targeting either SR-B1 or CD81 (Figure 1C) (Grove et al., 2007, 2017). CD81 blockade prevented entry of both WT and mutant equally - consistent with CD81 being an essential receptor (Figure 1B) - whereas the I438V A524T mutant was largely resistant to inhibition by anti-SR-B1. We corroborated this finding using BLT-4, a small molecule inhibitor of SR-B1 (Figure S3). Next, we used lentiviral vectors to over-express either receptor (Figure 1D). WT virus exhibited a strong response to increasing availability of SR-B1 and CD81, reaching a 3-4 fold enhancement relative to untreated cells. In contrast, I438V A524T HCV was less responsive than WT to increased CD81 availability, and was only modestly affected by SR-B1 over-expression. Finally, we directly evaluated E2-receptor interactions by measuring the binding of soluble E2 (sE2) to human SR-B1 or CD81 exogenously expressed on the surface of CHO cells (Figure 2E & F). I438V A524T sE2 exhibited altered receptor interactions, with >2 fold reduction in SR-B1 binding and >2 fold increase in CD81 interaction. In summary, HCV I438V A524T exhibits low dependency on SR-B1 and enhanced interactions with CD81. Of note, the reduction in SR-B1 binding by I438V A524T sE2 would suggest modulation of the SR-B1 binding site, HVR-1, even though neither of the mutated residues reside in this region.

**Fig. 2.**
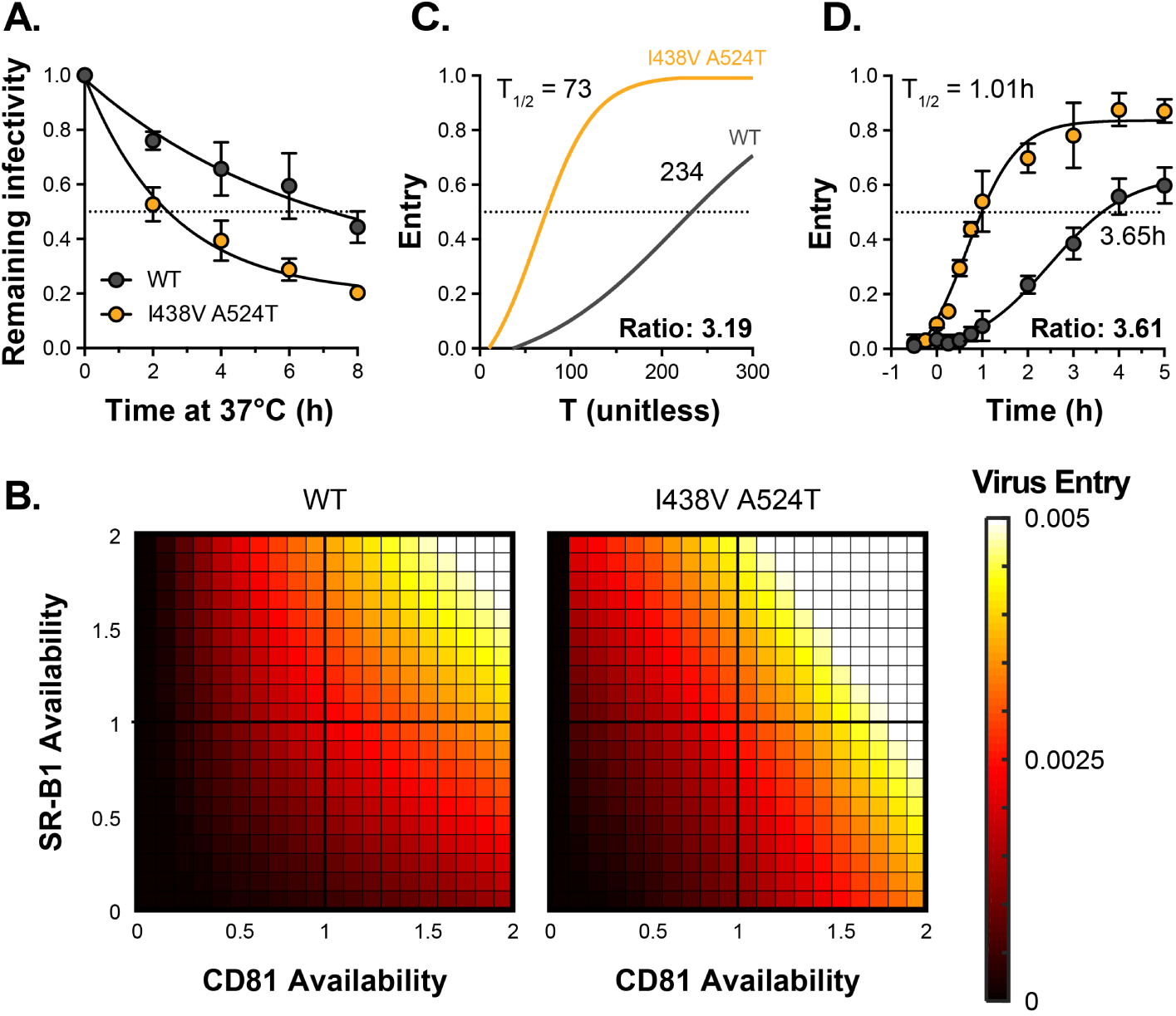
HCV I438V A524T is hyper-reactive. **A**. WT and I438V A524T HCVcc were incubated for 0-8 hours at 37°C before infection of Huh-7.5 cells. Remaining infectivity is expressed relative to t=0 time point. Data points represent the mean of two independent experiments, data was fitted using an exponential decay function. **B**. Mathematical modelling was used to predict the entry characteristics of WT and I438V A524T HCVcc (also see Figure S4). Heat maps display the probability of virus entry (as denoted in the key), for any given virus particle, upon varying availability of SR-B1 or CD81. Receptor availability is scaled relative to parental Huh-7.5 cells **C**. Kinetics of WT and mutant HCVcc entry, as predicted by mathematical modelling. The data is normalised to maximum entry. T represents uncalibrated time and, therefore, cannot be converted to real time, but relative differences can be estimated. Comparison of time to 50% entry suggests that I438V A524T HCVcc completes entry ∼ 3 times faster than WT. **D**. Kinetics of WT and mutant HCV entry were experimentally measured by synchronised infection of Huh-7.5 cells by HCVpp, followed by chase with a saturating inhibitory concentration of anti-CD81 mAb. Mutant HCVpp escaped the inhibitory effects of anti-CD81 ~ 3.5 times faster than WT. Data points represent the mean of three independent experiments. The data was fitted with a sigmoid function. In all plots error bars indicate standard error of the mean, fitted curves were confirmed to be significantly different (F-test, GraphPad Prism).

We recently developed a mathematical model to explore the mechanisms of HCV entry (Kalemera et al., 2019). This work supports the notion that E1E2 must acquire CD81 to proceed through the entry process and that this can be achieved via two routes: 1) SR-B1-mediated acquisition, where interaction with SR-B1 primes subsequent CD81 binding, or 2) intrinsic acquisition of CD81, without the necessity for prior engagement of SR-B1 (Figure S4A). The latter pathway accounts for residual infection by WT virus in the absence of SR-B1 (Figure 1B & C). Whilst this model is supported by published data (Augestad et al., 2020; Kalemera et al., 2019; Prentoe et al., 2019), the mechanistic basis of SR-B1 mediated priming has, thus far, been unclear. To recapitulate the physical reality of virus entry, the model includes a variable that reflects the intrinsic instability of HCV particles. Therefore, we compared the stability of WT and I438V A524T particles at 37°C. Unexpectedly, we found that mutant HCV was >3 fold less stable than WT virus (assessed by comparison of half-life, Figure 2A), this would suggest that the I438V A524T mutations increase the propensity for E1E2 to undergo spontaneous inactivation.

We integrated the measurements of receptor dependency and particle stability (Figure 1C-F, 2A) into our mathematical model, allowing us to compare the entry characteristics of WT and mutant HCV. This analysis predicts that the I438V A524T mutant exists in a hyper-reactive state wherein acquisition of CD81, via either route, is enhanced (SR-B1-mediated acquisition, in particular, being ~ 1000 fold more efficient, Figure S4B, parameter c_2_). Indeed, this would account for the ability of this mutant to tolerate reductions in SR-B1 availability (Figure 1B & C, S3). Moreover, down-stream entry events, encompassing cell surface translocation, endocytosis and fusion (Baktash et al., 2018), were predicted to occur at ~ 6-fold higher rate (parameter e, Figure S4B). The model also allows exploration of the stoichiometry of HCV-CD81 interaction. To this end, fitting the combined WT and mutant data indicated that the acquisition of two CD81 molecules is sufficient for a HCV particle to proceed along the entry pathway (Figure S4C), and this was the case for both WT and mutant. This value is somewhat lower than our previous estimate; however, in our current study we have fitted the model with more data and achieved a distinct, and decisive, peak in likelihood, unlike our prior estimate which had a very broad range of possible stoichiometries (Kalemera et.al. 2019).

We used the model to make further predictions. First, we estimated the entry of WT and mutant HCV upon varying availabilities of SR-B1 and CD81. Here the I438V A524T mutant exhibited increased entry efficiency over a greater range of receptor densities (Figure 2B). An expected consequence of this enhanced entry efficiency is an increase in the kinetics of entry. We therefore used the model to estimate the relative speed at which WT and mutant HCV complete entry (Figure 2C), and in parallel made measurements of entry kinetics in vitro using synchronised HCV psedudoparticle (HCVpp) infection (Figure 2D). The model predicted that mutant HCV enters >3 fold faster than WT. This estimate was in excellent agreement with the experimentally measured values, corroborating our modelling approach.

In summary, HCV is capable of evolving to optimise virus entry. It achieves this by adopting a hyper-reactive state, which exhibits low SR-B1 dependency, rapid acquisition of CD81 and increases in downstream entry events. However, the benefits of hyper-reactivity are somewhat offset by a decrease in stability. This likely reflects a greater propensity for E1E2 to undergo spontaneous and irreversible inactivation, possibly through premature triggering of its fusion activity.

### Hyper-reactive HCV is acutely sensitive to all neutralising antibodies

The majority of HCV^+^ patients experience chronic life-long infection with persistently high viral loads. To achieve this, HCV must resist the E1E2-specific nAbs that arise in most individuals. Failure of HCV to evade and/or escape nAbs has been linked with viral clearance and understanding the molecular mechanisms of HCV antibody resistance is likely to inform ongoing HCV vaccine development (Keck et al., 2018; Kinchen et al., 2018, 2019).

To evaluate the capacity of HCVcc to resist nAbs we measured sensitivity to chronic HCV patient immunoglobulins (IgGs). WT HCV was highly resistant to patient nAbs (Figure 3A); the neutralisation curve following a log-linear relationship, such that successive 10-fold increases in IgG concentration yielded only a modest increase in neutralisation. In stark contrast, the hyper-reactive HCV I438V A524T mutant was acutely sensitive to patient IgG, reaching complete neutralisation even at low concentrations of nAb.

Intense investigation of anti-HCV nAb responses, by others, has provided a detailed understanding of the major antigenic targets in E2 (Figure 3B); for example many potent nAbs target one or other component of the CD81 binding site (AS412, Front Layer and CD81 Binding Loop) (Tzarum et al., 2018). We measured the sensitivity of WT and mutant HCV to a panel of mAbs targeting a range of these antigenic targets (Figure 3C & S5A) (Bailey et al., 2017; Giang et al., 2012; Pierce et al., 2016; Sabo et al., 2011). Without exception, WT virus resisted mAb neutralisation whereas I438V A524T HCV was potently inhibited; this is best illustrated by examining the change in IC50 values across the panel of nAbs, which demonstrates a ~ 20 fold increase in nAb sensitivity (Figure 3D). We also assessed the ability of soluble CD81 EC2 to neutralise HCVcc (Figure 3G) and observed the same pattern of inhibition. To evaluate whether this global shift in nAb sensitivity reflected changes in antibody binding we measured nAb interactions with sE2 by ELISA (Figure 3E & S5B). Without exception, mAbs bound equally to WT and I438V A524T E2, this would suggest that WT and mutant E2 are antigenically equivalent.

**Fig. 3.**
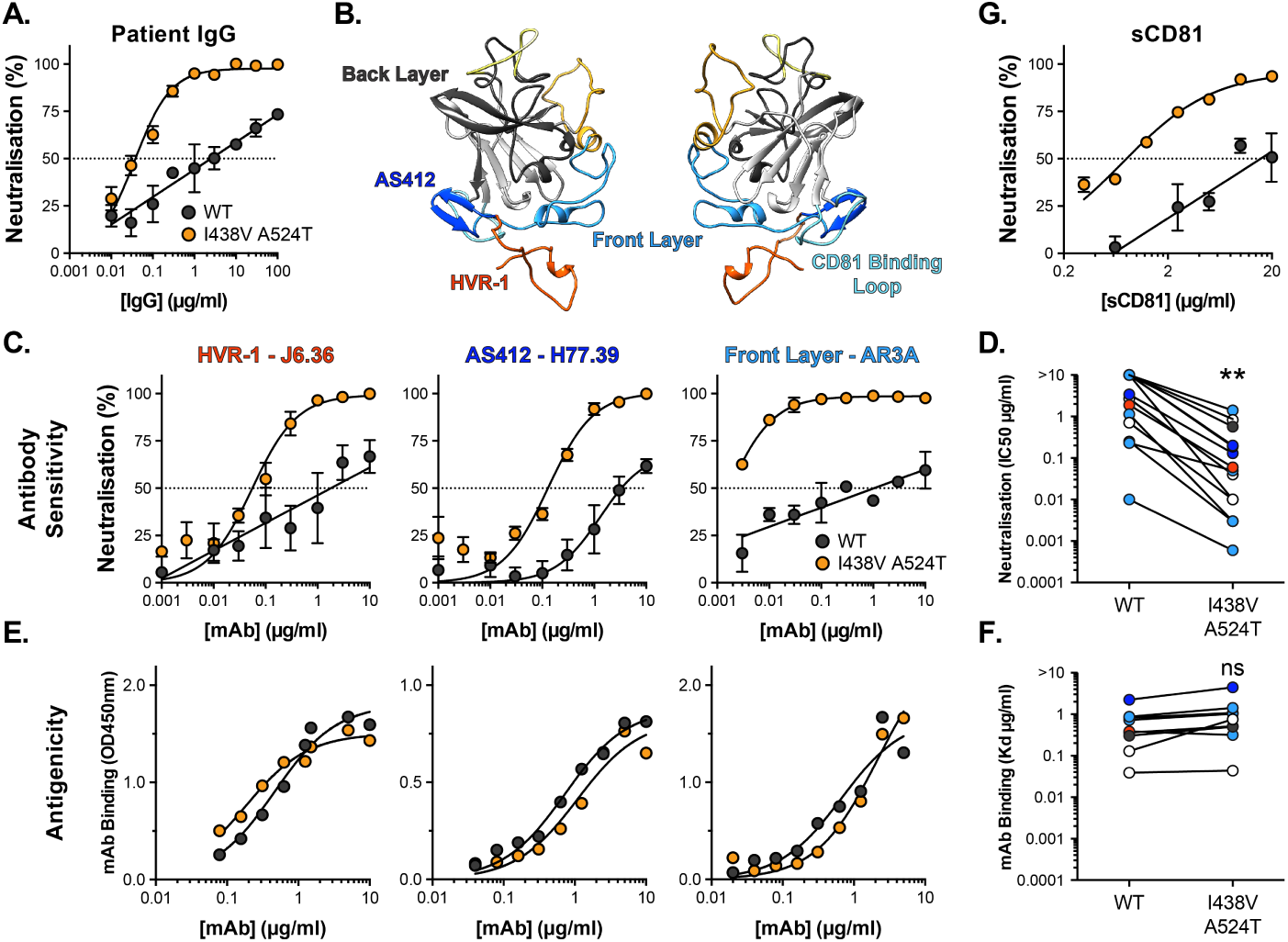
Hyper-reactive HCV is acutely sensitive to all neutralising antibodies. **A**. Neutralisation of WT and I438V A524T HCVcc by a serial dilution of HCV patient IgG. **B**. Molecular cartoon of major antigenic sites targeted by nAbs. **C**. Neutralisation curves for three representative mAbs targeting distinct sites, the mAb name and specificity are provided and color coded to match B. **D**. 50% inhibitory concentrations of 14 mAbs, data points are color coded according to their antigenic target, aster-isks indicate statistical significance (T-test, Graph-Pad Prism). **E**. Binding of mAbs to WT and I438V A524T E2 assessed by ELISA; mAbs matched to neutralisation data. **F**. Estimated dissociation constants for 9 mabs, data points color coded as above, there is no significant difference between WT and mutant (T-test, GraphPad Prism). **G**. HCVcc neutralisation by soluble CD81 EC2. For all neutralisation curves, data points represent the mean of n=2 or 3 independent experiments. I438V A524T data were best fitted by a hyperbola function (GraphPad Prism), WT data were best fitted by a semi-log function (except for H77.39). For ELISA data, one representative experiment is shown, with data fitted with a hyperbola function. In all plots error bars indicate standard error of the mean. Estimates of IC50s and Kd were obtained from the fitted curves.

In summary, the I438V A524T mutant exhibits acute sensitivity to antibody-mediated neutralisation without intrinsic changes to the antigenicity of E2. This would suggest that whilst nAb binding remains unaltered, the consequences of antibody binding are dramatically changed. This phenotype likely reflects the hyper-reactive state of the I438V A524T mutant; much like incubation at 37°C (Figure 2A), nAb interactions may trigger irreversible inactivation of E1E2.

### Hyper-reactive HCV exhibits stabilisation of HVR-1

The antigenic similarity of WT and I438V A524T E2 would suggest there is no gross conformational change associated with the hyper-reactive phenotype. However, when subjected to limited proteolysis I438V A524T E2 undergoes cleavage more rapidly than WT E2 (Figure S6), this suggests that subtle structural differences may underpin hyper-reactivity. We have previously used molecular dynamic simulations (MD) to explore the conformational landscape of E2, finding that flexibility and disorder are conserved features of E2, consistent with other reports (Balasco et al., 2018; Kong et al., 2016; Meola et al., 2015; Stejskal et al., 2020; Ströh et al., 2018; Vasiliauskaite et al., 2017). In particular, we discovered that HVR-1 (which contains the SR-B1 binding site), is a putative intrinsically disordered peptide tail. Therefore we used MD to examine the motion of E2, performing five independent 1*µ*s simulations of WT and I438V A524T E2 ectodomain. The overall dynamics of either E2 was similar, as reflected in their root mean square fluctuation profiles (RMSF; Figure 4C). However, the HVR-1 tail of I438V A524T E2 exhibited consistent stabilisation (Figure 4C, S7, S8), this is best illustrated by root mean square deviation (RMSD), which captures motion over time (Fig. 4A, 4B, S7 & S8), and RMSF of HVR-1 (Figure 4C, inset).

**Fig. 4.**
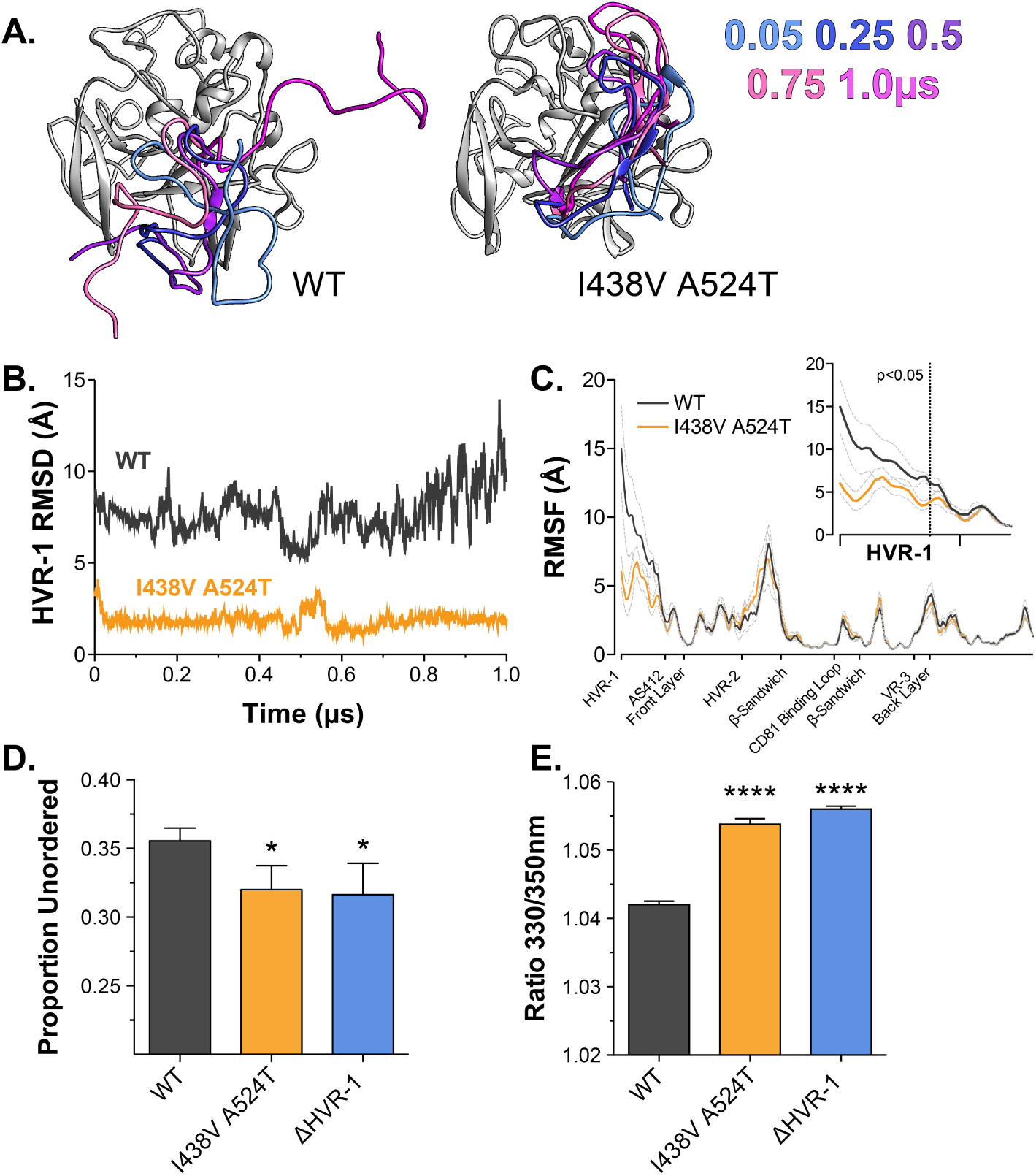
Hyper-reactive HCV exhibits stabilisation of HVR-1. The conformational dynamics of WT and I438V A524T E2 ectodomain were explored by 5 independent 1*µ*s MD simulations. **A**. Images summarising two representative simulations; HVR-1 is color coded according to time (as shown in key), the remainder of E2 is shown in grey for the t=0.05*µ*s frame only. **B**. Root mean square deviation (RMSD) of HVR-1 for the simulations shown in A. **C**. Average root mean square fluctuation (RMSF) for WT and I438V A524T E2, from five independent experiments, error bars indicate standard error of the mean. X-axis denotes regions of E2. Inset provides a zoom of the data for HVR-1, RMSF values to the left of the dashed line reach statistical significance (ANOVA, GraphPad Prism). Summaries of all MD simulations are provided in Figure S7 and S8. **D**. Estimation of un-ordered protein content, by circular dichroism spectroscopy, for WT, I438V A524T and ΔHVR-1 sE2. Data represent the mean of three independent measurements. **E**. Intrinsic fluorescence ratio (330nm over 350nm), measured by nano differential scanning fluorimetry, for WT, I438V A524T and ΔHVR-1 sE2 at 37°C. Data represent the mean of three independent measurements. Asterisks denote statistical significance (ANOVA, GraphPad Prism).

Given these data, we reasoned that I438V A524T E2 may share biophysical characteristics with E2 lacking HVR-1 (ΔHVR-1, in which HVR-1 is genetically deleted). We first analysed sE2 by circular dichroism spectroscopy (CD), which provides a low-resolution ensemble measurement of protein structure (Figure S9). Here, I438V A524T and ΔHVR-1 E2 were biophysically indistinguishable from one another but distinct from WT E2. In particular, estimations of the structural composition of I438V A524T and ΔHVR-1 E2 indicated a significant reduction of unordered components (Figure 4D); this is broadly consistent with our MD experiments. We also assessed sE2 by nano differential scanning fluorimetry (nanoDSF), which exploits the changes in intrinsic protein fluorescence upon solvent exposure of tryptophan residues to provide a surrogate measure of protein folding and to determine melting temperature (Figure S10). Consistent with other reports, the apparent melting temperature of sE2 was high (>80°C; Figure S10B) (Kong et al., 2016), this likely represents unfolding of the stable globular core of E2 (i.e. *β*-sandwich; Figure S1C), and, in this respect, we observed no differences between each E2. However, at lower, physiological, temperatures the intrinsic fluorescence ratio of I438V A524T and ΔHVR-1 E2 were higher than WT (Figure S10A & 4E), consistent with mutant and ΔHVR-1 E2 being biophysically distinct from WT. Notably, HVR-1 is directly upstream of a highly conserved tryptophan residue in AS412 that has been previously demonstrated to be important for CD81 binding (W420, (Cowton et al., 2016; Owsianka et al., 2006)).

In summary, MD simulations suggest that E1E2 hyper-reactivity is associated with tuning of HVR-1 dynamics. This notion is supported by the biophysical similarity of mutant and ΔHVR-1 E2. Therefore, HVR-1 dynamics provide a potential molecular mechanism that links E1E2 reactivity, SR-B1 receptor dependency, entry efficiency and nAb sensitivity.

### Hypervariable region-1 is an entropic safety catch

Based on our data, we hypothesise that HVR-1 exerts an autoinhibitory effect on E1E2 that is dependent on its conformational entropy and/or dynamics. Furthermore, during virus entry, engagement of SR-B1 will, necessarily, constrain HVR-1, reduce conformational entropy (Smyda and Harvey, 2012) and turn off autoinhibition. Therefore, HVR-1 is analogous to a safety catch on a firearm, that suppresses the reactivity of E1E2; SR-B1 turns off the safety catch at the cell surface to prime virus entry. In the hyper-reactive I438V A524T mutant, pre-stabilisation of HVR-1 turns off the safety catch on free virions, reducing their dependency for SR-B1 and enhancing entry efficiency, but rendering E1E2 more prone to spontaneous triggering and inactivation.

Tuning of protein function by disordered peptide tails have been described in other systems (Uversky, 2013); in these cases the tail can generate an entropic force which acts upon the rest of the protein (Keul et al., 2018). We examined this possibility in E1E2 using dynamic cross correlation (DCC), which provides a measure of correlated/coordinated motion within MD simulations. DCC analysis of WT E2 suggests that motions within HVR-1 are transmitted throughout the protein (Figure S11), consistent with an exerted force. This likely occurs through allosteric linkages, via disulphide bonds (Figure S11, boxed regions), from the N-terminal region (HVR-1 & AS412) through to the central *β*-sandwich and C-terminal Back Layer. In the simulations of I438V A524T E2, HVR-1 is stabilised (Figure 4) and these correlated motions are absent (Figure S11), suggesting the lack of an entropic force. This provides a potential molecular mechanism by which HVR-1 dynamics can influence the entire protein.

Given these observations, we reasoned that genetic removal of HVR-1 would prevent this entropic force, turn off the safety catch and confer a hyper-reactive phenotype. We characterised the reactivity of WT, I438V A524T and ΔHVR-1 HCVcc by measuring infectivity, receptor dependency, thermal stability and neutralisation sensitivity (Figure 5). In each case, ΔHVR-1 HCVcc exhibited a hyper-reactive phenotype above and beyond that of the I438V A524T mutant; ΔHVR-1 HCVcc has very high infectious titres, low SR-B1 dependency, thermal instability and acute sensitivity to patient IgG. These data are completely consistent with the entropic safety catch model.

**Fig. 5.**
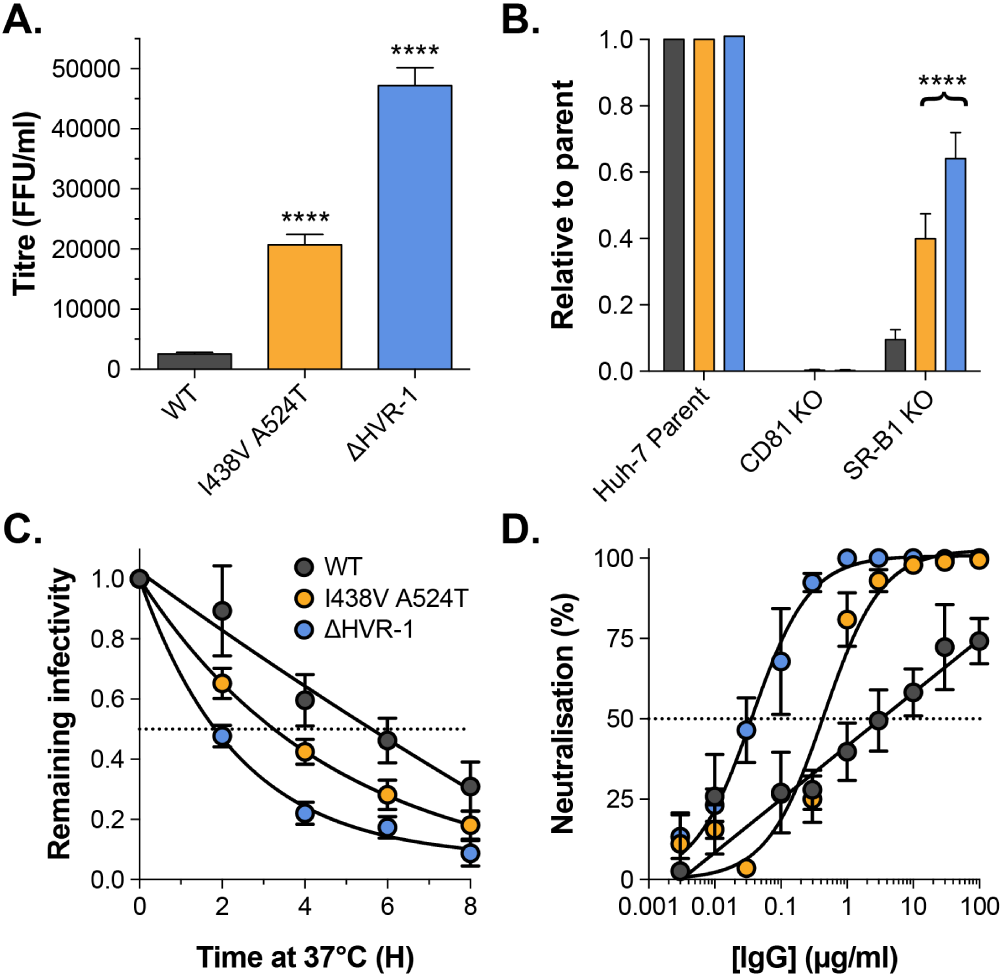
Removal of HVR-1 renders HCV hyper-reactive. **A**. Mean infectivities of WT, I438V A524T and ΔHVR-1 HCVcc; foci forming units are corrected for input particle numbers **B**. HCVcc infection of parental Huh-7 cells or those CRISPR/Cas9 edited to prevent expression of CD81 or SR-B1. Data is expressed relative to parental cells, mean of three independent experiments. **C**. Stability of HCVcc at 37°C, data points represent the mean of three independent experiments, data was fitted using an exponential decay function. **D**. Neutralisation of HCVcc by patient IgG, data points represent the mean of three independent experiments. Data was fitted with a hyperbola function (I438V A524T and ΔHVR-1) or semilog function (WT). In each plot error bars indicate standard error of the mean; asterisks denote statistical significance (ANOVA, Graphpad prism); all curves determined to be statistically significant (F-test, Graphpad prism).

### Antibody neutralisation prevents the selection of hyper-reactive mutants

Given the high infectivity of I438V A524T and ΔHVR-1 HCVcc, there remains a question over the advantages of evolving a mechanism to suppress E1E2 activity. The likely explanation is that in chronic infection nAbs necessitate tight control over E1E2 reactivity and that antibody selection prevents the emergence of hyper-reactive mutants. To test this we performed new culture adaptation experiments in the presence and absence of antibody selection by patient IgG, and measured the infectivity and antibody sensitivity of the resultant virus populations (Figure 6A & 6B). The antibody selected culture exhibited low infectivity and high resistance to neutralisation, much like WT HCVcc. In contrast, the culture without selection developed a hyper-reactive phenotype; having increased infectivity and high sensitivity to neutralisation.

**Fig. 6.**
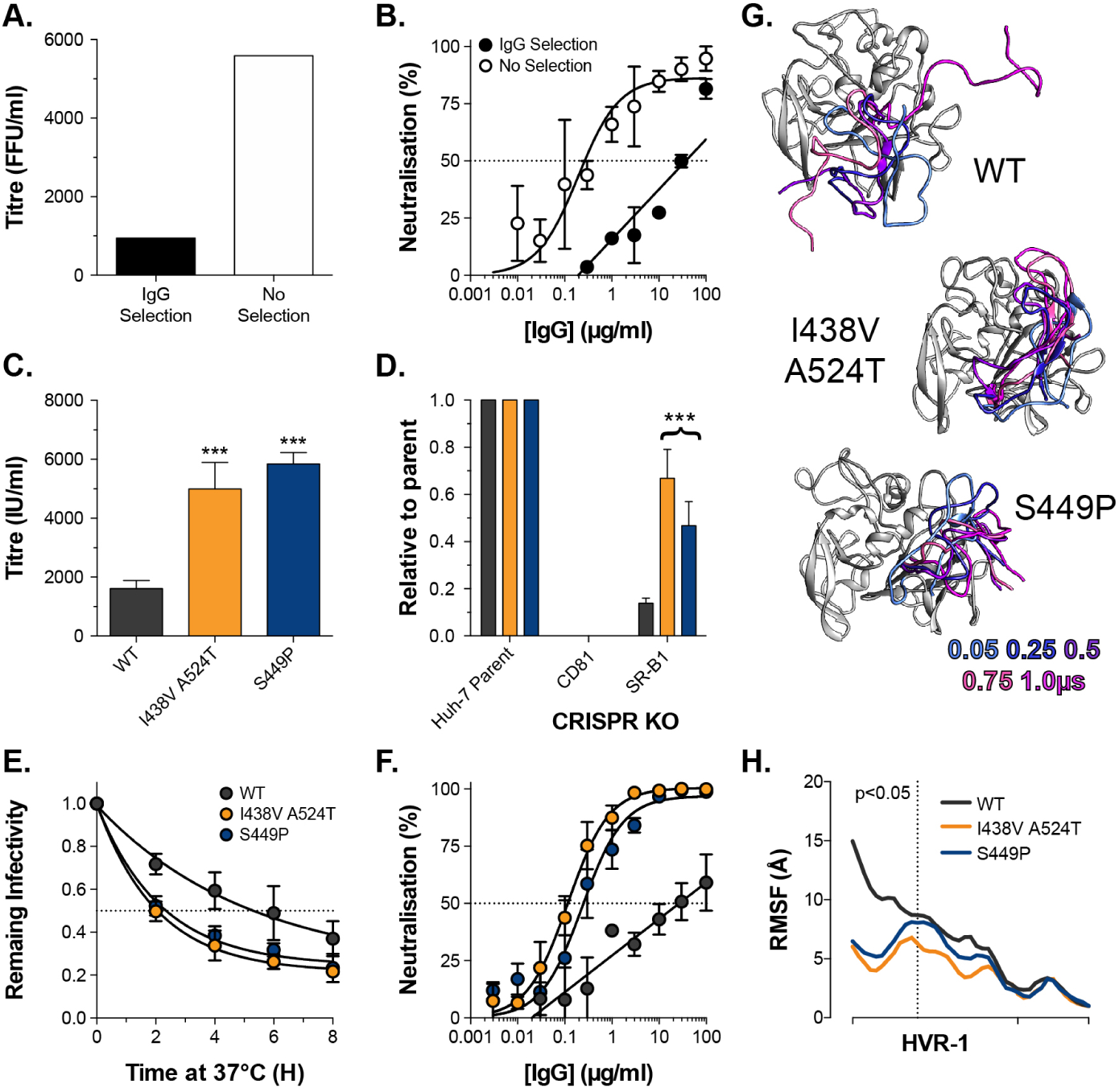
Emergence of hyper-reactive HCVcc in the absence of antibody selection. J6/JFH HCVcc was propagated in Huh-7.5 cells with and without antibody selection by HCV^+^ patient IgG. **A**. Infectious titre of HCVcc after propagation with and without antibody selection. **B**. Patient IgG neutralisation of HCVcc cultured with and without antibody selection, data points represent the mean of three technical repeats. **C**. Infectivity of WT, I438V A524T and S449P HCVcc. **D**. Receptor dependency of HCVcc, measured as in Figure 5B. **E**. Thermal stability of HCVcc, measured as in Figure 5C. **F**. Neutralisation sensitivity of HCVcc, measured as in Figure 5D. **G**. Images summarising representative MD simulations of WT, I438V A524T and S449P E2 ectodomain; HVR-1 is color coded by time, as denoted in the key, the remainder of E2 is shown in grey for t=0.05*µ*s only. **H**. Average RMSF of WT, I438V A524T and S449P HVR-1; values to the left of the dashed line reach statistical significance for both mutants compared to WT (ANOVA, GraphPad Prism).

NGS analysis of the culture without selection revealed a mixed population of WT and mutant viruses (not shown); this included an S449P mutant that was also identified at early time points in the original culture experiment (Figure S1C). Therefore, we used reverse genetics to introduce this single mutation and fully characterised the resultant virus. In every respect, S449P HCVcc exhibited a hyper-reactive phenotype that was indistinguishable from I438V A524T HCVcc (Figure 6C-F). Moreover, MD simulations of S449P E2 demonstrated stabilisation of HVR-1 to the same extent as I438V A524T (Figure 6G-H, S12). These data strongly support our HVR-1 safety catch model and suggest that emergence of hyper-reactive HCV can only occur in the absence of antibody selection. Therefore, the HVR-1 safety catch likely represents an important mechanism for maintaining HCV resistance to neutralising antibodies.

## Discussion

Here, we demonstrate that HVR-1 of HCV E2 is an autoinhibitory safety catch that regulates E1E2 reactivity, entry efficiency and antibody resistance. Our model suggests that, by constraining HVR-1, E2-SR-B1 interactions turn off the safety catch at the cell surface. Alternatively, HCV can evolve to pre-constrain HVR-1, therefore desuppressing E1E2 reactivity, however, this evolutionary pathway is blocked by neutralising antibody selection.

This model reconciles over a decade of research into HCV entry. It has long been established that HVR-1 deletion renders HCV sensitive to neutralising antibodies (Bankwitz et al., 2010, 2014; Prentoe et al., 2011, 2014, 2016). This originally led to the hypothesis that HVR-1 acts as a shield to occlude neutralising epitopes and the CD81 binding site. However, it has since become clear that the global nature of nAb sensitisation is not easily explained by a simple shielding model (Augestad et al., 2020; Prentoe et al., 2019). Moreover, many studies have noted that substitutions in E2, many arising during in vitro culture, confer altered SR-B1 dependency and sensitivity to neutralisation (Bitzegeio et al., 2010; Dhillon et al., 2010; Grove et al., 2008; Keck et al., 2009; Koutsoudakis et al., 2012; Lavie et al., 2014; Song et al., 2012). Finally, various studies have provided evidence that initial interactions with SR-B1 may act to prime subsequent events in HCV entry (Kalemera et al., 2019; Prentoe and Bukh, 2018). Indeed, Augestad et. al. have recently suggested that a conformational opening of E1E2 may be regulated by SR-B1-HVR-1 interactions (Augestad et al., 2020). The HVR-1 safety catch model sufficiently rationalises all of the above observations and, therefore, offers mechanistic clarity on the early events of HCV entry

A unique feature of HCV particles is their association with host-derived lipoprotein components, resulting in so-called lipo-viro-particles. Lipoprotein receptors and cholesterol transporters have been implicated in HCV entry (Ding et al., 2014; Lindenbach and Rice, 2013); indeed, SR-B1 is a high-density lipoprotein receptor and it has been suggested that it may interact with host apolipoproteins on HCV virions (Maillard et al., 2006). We have not considered the contribution made by lipoprotein components in our studies, however, the hyper-reactive phenotype is recapitulated in HCV pseudoparticles (Figure S13), which are lacking any lipoprotein components. Therefore, it is not necessary to invoke lipoprotein involvement in our model. It is most likely that lipoprotein components contribute to cell surface attachment prior to E2-receptor engagement. Indeed, ΔHVR-1 HCVcc is modestly inhibited by SR-B1 KO (Figure 5), even though E2-SR-B1 interactions are not possible in this virus (Scarselli et al., 2002); this may reflect the absence of particle tethering to the cell surface through SR-B1-apolipoprotein interactions.

An implication of our model is that HCV entry efficiency and nAb resistance are opposing selection pressures that can be balanced through tuning of HVR-1 dynamics. Various studies have identified patient derived E2 polymorphisms (including position 438) that alter antibody sensitivity, receptor usage and infectivity; these observations are consistent with changes in E1E2 reactivity (Bailey et al., 2015; El-Diwany et al., 2017; Fofana et al., 2012; Keck et al., 2009). This would suggest that HCV can trade reductions in entry efficiency for increases in nAb resistance, and vice versa. This may be particularly advantageous over the course of natural infection. During the early acute phase of infection the Ab response has not yet matured, this may allow HCV evolution to turn off the safety catch in favour of enhanced virus entry, allowing the infection to be established and consolidated. Whereas in later chronic infection the emergent nAb response may force HCV to turn on the safety catch and restore nAb resistance. Therefore, the HVR-1 safety catch likely represents an adaptation to the ongoing immune assault in chronic infection.

HVR-1 is, itself, an immunodominant target for antibodies; however, anti-HVR-1 responses are rapidly evaded through mutational escape (von Hahn et al., 2007). Our model would suggest that disorder and conformational entropy are critical to HVR-1 function. This feature likely underpins its ability to tolerate extensive variation. Antigenic variation, driven by nAb selection, is detrimental to other functionally important regions of E2 (Keck et al., 2009; Kinchen et al., 2018). On the contrary, for HVR-1 continuous antigenic drift likely ensures that disorder is maintained. This provides a potential feedback system where nAb selection promotes conformational entropy and keeps the safety catch mechanism engaged.

A complete structure-to-function understanding of E1E2 will likely reveal a novel viral fusion mechanism and guide the rational design of HCV immunogens. An appreciation of the importance of HVR-1 in regulating E1E2 reactivity will guide future investigations in these areas. For example, many structural studies of E2 have been performed on ΔHVR-1 constructs; our data suggest that without the HVR-1 safety catch mechanism, E1E2 is more likely to undergo spontaneous inactivation. Therefore, HVR-1 truncation may hinder attempts to solve the pre-fusion structure of E1E2. Regarding vaccine development, a recent study used ΔHVR-1 E1E2 as an immunogen (guided by the notion that HVR-1 acts as an epitope shield) and found it to be inferior to WT E1E2 (Law et al., 2018); this may, again, reflect the spontaneous transition of ΔHVR-1 E1E2 to a non-functional state. Vaccine development in other fields suggests that locking entry proteins in the pre-fusion state may be critical for eliciting potent nAb responses (Crank et al., 2019; Hsieh et al., 2020; Rey and Lok, 2018; Sanders et al., 2015; de Taeye et al., 2015). Thus, engineering the HVR-1 safety catch to improve E1E2 stability may be a viable strategy for HCV immunogen design.

We have discovered that HVR-1 acts as a safety catch and have identified potential allosteric linkage between HVR-1 and distant regions of E2 (Figure S11), however, we do not yet have a complete molecular understanding of this autoinhibitory mechanism. Our future investigations will aim to understand how E2-SR-B1 interactions toggle the HVR-1 safety catch and identify the structural/dynamical consequences for the E1E2 complex. For example, Augestad et. al. have suggested that SR-B1 may trigger a conformational opening of E1E2, associated with changes in AS412 (a functional epitope directly downstream of HVR-1); although the structure of this open state, and how it relates to E1E2 reactivity, remains unclear (Augestad et al., 2020). Alternatively, Tzarum et. al. recently identified a conformational switch in the front layer of E2 that may be important for CD81/antibody engagement (Tzarum et al., 2020). Might these events be regulated by the safety catch?

Notably, the hyper-reactive phenotype, identified here (receptor independency, reduced stability, nAb sensitivity), shares striking parallels with a hyper-reactive state adopted by HIV Env (Haim et al., 2011). This reinforces the message that all viral entry proteins perform similar thermodynamic balancing acts. Therefore, might other viruses harness structural disorder to regulate entry? Intrinsically disordered tails have been commonly described in eukaryotes, where they regulate protein function and have autoinhibitory activity (Trudeau et al., 2013; Uversky, 2013). The apparent tunability of disordered protein tails is likely to synergise with the genetic plasticity of viruses. Indeed, the entropic safety catch may yet prove to be a common strategy for navigating the thermodynamic landscapes of virus entry and antibody evasion.

## Materials and methods

### Cell cultures

Huh-7.5 cells were acquired from Apath LLC. CRISPR Cas9 receptor KO Huh-7 cells (Yamamoto et al., 2016), and parental Huh-7 cells, were generously provided by Yoshiharu Matsuura (Osaka University, Japan). CHO and HEK293T were acquired from the American Type Culture Collection. All cells were grown at 37°C in Dulbecco’s Modified Eagle Medium (DMEM) supplemented with 10% foetal calf serum (FCS), penicillin & streptomycin and non-essential amino acids.

### Antibodies

Mouse Anti-NS5 mAb (S38) and anti-Cd81 mAb (2.131) were a gift from Prof. Jane McKeating (University of Oxford). Rabbit Anti-SR-B1 serum was provided by Dr. Thierry Huby (INSERM, Paris). Mouse anti-E2 mAbs J6.36 and H77.39 were a gift from Michael Diamond (Washington University). Human anti-E2 mAbs were kindly provided by Dr. Mansun Law (SCRIPPS, La Jolla) (AR2A, AR3A, AR3C, AR4A, AR5A), Dr. Steven Foung (Stanford) (HC33.1.53, HC84.26, CBH-7, HC1, CBH-4B, CBH-7, CBH-23), and Prof. James Crowe (Vanderbilt University) (HepC3, HepC43). StrepMAB-classic was purchased from IBA Lifesciences (Göttingen, Germany).

### Isolation of patient IgG

Blood samples were collected from HCV+ patients under ethical approval: “Characterising and modifying immune responses in chronic viral hepatitis”; IRAS Number 43993; REC number 11/LO/0421. Extracted serum was heat inactivated, filtered and diluted 1:1 in PBS. Total IgG was captured using a HiTrap protein G column (Cytiva, MA, USA), eluted in pH2.7 glycine buffer and buffer exchanged into PBS. For experiments, batches of pooled IgG were created by equimolar combination of IgG from two patient samples.

### Production of HCVcc

Plasmid encoding cell-culture proficient full-length J6/JFH-1 (acquired from Apath LLC) was used as a template for the in vitro production of infectious HCV RNA (Lindenbach et al., 2005). To initiate infection, viral RNA was electroporated into Huh-7.5 cells using a BTX830 (Harvard Instruments, Cambridge, UK). From 3-7 days post electroporation, cell culture supernatants containing infectious J6/JFH-1 HCVcc were harvested every 3-5 hours. Short harvest times limit the opportunity for virus decay thereby preventing the accumulation of non-infectious particles. To ensure maximum reproducibility between experiments, a standardized stock of experimental virus was generated by pooling the harvested supernatants.

### HCVcc adaptation experiments

In the initial adaptation experiment (Figure 1) a continuous culture of J6/JFH HCVcc was established in Huh-7.5 cells. Infected cells were passaged, twice weekly, with an excess of uninfected target cells added whenever the culture reached 90-100% infection. This proceeded for 20 weeks. For the adaptation experiments with and without IgG selection (Figure 6) we adopted a serial passage strategy. Huh-7.5 cells were electroporated with the J6/JFH genome and the next day medium was replaced with DMEM 3% FCS plus or minus HCV^+^ patient IgG at 100*µ*g/ml. Once 90-100% of cells were determined to be infected, the culture supernatant was harvested, clarified by centrifugation, and used to infect fresh Huh-7.5 cells. Typically, this was done every week. In either case (continuous culture or passage) cell culture supernatants were frozen at −20°C at regular intervals for analysis by NGS.

### HCVcc infections

Huh-7.5 and receptor KO Huh-7 cells were seeded at 1.5 x 10^4^ cells per well of a 96 well plate 24 hours prior to the experiment. To quantify infectious titres, cells were challenged with a two-fold serial dilution of virus stock (1/4 to 1/64) in DMEM 3% FCS (infection medium). Cells were incubated with viral supernatants for five hours before the cells were subsequently washed, and fresh medium was added. Infections were allowed to proceed for 48 (Huh-7.5 cells) or 72 (Huh-7 lines) hours before reading out. HCVcc replication was quantified by fluorescence microscopy. Cells were fixed with 100% methanol and stained for viral NS5A protein with mAb S38. HCV foci-forming units (FFU) were quantified by manual counting, or percentage infection determined by automated analysis of fluorescence micrographs (Culley et al., 2016).

### Neutralisation/inhibition assays

Huh-7.5 cells were seeded, as above. For neutralisation assays, virus was preincubated with a dilution series of mAb or soluble CD81 EC2 for 1 hour at 37C prior to infection. For anti-receptor blockade experiments, Huh-7.5 cells, seeded for infection, were pre-incubated at 37°C with 50 *µ*l DMEM 3% FCS containing a serial dilution of either rabbit-SR-B1 serum or mouse anti-CD81 mAb 2.131. One hour later, wells were challenged with virus. In each case the infections were processed as described above.

### Next generation sequencing and analysis

RNA was extracted from cell culture supernatants containing HCVcc by a BioRobot MDx instrument using QIAamp Virus BioRobot MDx Kits. Extracted RNA samples were amplified as described (Aisyah et al., 2019) processed locally within the UCL Hospital Virology laboratories for PCR library preparation and Next Generation Sequencing using Illumina MiSeq equipment (Manso et al., 2020). Sequences were trimmed of adaptors and low quality reads using Trimmomatic V. 0.33 (Bolger et al., 2014). The quality of the sequence files was then assessed using the FastQC program. The resulting FASTQ files were then aligned to the indexed reference J6/JFH HCVcc genome using the BWA-MEM algorithm (Burrows Wheeler Aligner (Li and Durbin, 2009)), converted into Sequence Alignment Map (SAM) files, which were further compressed into BAM files (binary versions of SAM files), sorted by reference coordinates and indexed using SAMtools. The duplicate sequences were then removed by Picard Tools and indexed again using SAMtools. Base-calling for each position in the genome was extracted from the indexed file. Positions of interest were identified as those with at least 1000 reads available with variance in nucleotide base composition of *≥* 5%.

### Production of lentiviral vectors and receptor over-expression

To generate lentiviral vectors, HEK293T cells were transfected with three plasmids: an HIV packaging construct (pCMV-dR8.91), VSV-G envelope plasmid (pMD2.G) and a transfer plasmid encoding GFP and SR-B1 or CD81, expressed from separate promoters (pDual SR-B1 or CD81, available from Addgene: https://www.addgene.org/Joe_Grove/). Supernatants containing viral vectors were collected at 48 and 72 hours post-transfection. At least 96 hours before an experiment, Huh-7.5 were transduced with lentivirus vectors diluted in complete medium and 24 hours prior to study the cells were seeded into a 96 well plate for infection, as described above.

### Production of HCV pseudoparticles

To generate HCVpp, HEK293T^CD81KO^ cells (Kalemera et al., 2020) were co-transfected with three plasmids: an HIV packaging construct (pCMV-dR8.91), a luciferase reporter plasmid (CSLW) and an expression vector encoding the appropriate HCV glycoprotein. Supernatants containing HCVpp were collected at 48- and 72-hours post-transfection.

### Entry kinetics assay

Huh-7.5 cells were seeded for infection, as above. HCVpp were preincubated with magnetic nanoparticles (ViroMag, OZ Biosciences, France) for 15 min, added to Huh-7.5 cells and the plate was placed on a Super Magnetic Plate (OZ Biosciences, France) for 15 min at 37°C to synchronise infection (Haim et al., 2005, 2009). The synchronous infection was chased with a saturating receptor blockade by adding 3*µ*g/ml anti-CD81 mAb 2.131 at the indicated time points. Infection was assayed after 72 hours using the SteadyGlo reagent kit and a GloMax luminometer (Promega, USA).

### Production of sE2

Soluble E2 comprises residues 384-661 of the HCV genome, flanked by an N-terminal tissue plasminogen activator signal sequence and a C-terminal Twin-Strep-tag. HEK293T^CD81KO^ cells (Kalemera et al., 2020) were transduced with lentivirus encoding sE2. Cell culture media, containing sE2, were harvested every 24 hours for up to 6 weeks. The harvested supernatants were frozen immediately at −80°C. High purity monomeric sE2 was generated by sequential affinity purification using StrepTactin-XT columns (IBA Lifesciences, Göttingen, Germany) and size-exclusion chromatography using the HiPrep™ 16/60 Sephacryl® S-200 HR gel filtration column (Cytiva, MA, USA).

### Soluble E2 binding assay

The sE2 binding assay has been described elsewhere (Kalemera et al., 2019). Briefly, a single-cell suspension of 5×10^5^ CHO cells transduced to express human SR-B1 or CD81 (protocol for receptor lentiviral production described above). Cells were preincubated in ‘traffic stop’ buffer, PBS + 1% bovine serum albumin and 0.01% sodium azide; the addition of NaN_3_ depletes cellular ATP pools, consequently preventing active processes including receptor internalisation. All subsequent steps are performed in traffic stop buffer. Cells were pelleted and then resuspended in a serial dilution of sE2. Following a 1 hour incubation at 37°C, cells were washed twice and incubated with 3*µ*g/ml StrepMAB-classic (for CHO SR-B1) or J6.36 (for CHO CD81) followed by an anti-mouse Alexa Fluor 647 secondary. After a final wash, the cells were fixed in 1% formaldehyde and analysed by flow cytometry. To measure cell surface expression of human receptors post-transduction, cells were stained for SR-B1 or CD81 and signal was detected using an anti-rabbit or anti-mouse Alexa Fluor 647 secondary, respectively.

### ELISA

Purified Strep-II-tagged sE2 monomers (1.0 *µ*g/mL diluted in TRIS buffered saline, TBS) were added to 96-well Strep-TactinXT-coated plates (IBA Lifesciences, Göttingen, Germany) for 2 hours at room temperature. Plates were washed with TBS twice before incubating with serially diluted mAbs in casein blocking buffer (Thermo Fisher Scientific) for 90 min. After three washes with TBS, well were incubated with a 1:3000 dilution of HRP-labelled goat anti-human/mouse IgG in casein blocking buffer for 45 min. After washing the plates five times with TBS+0.05% Tween-20, plates were developed by adding develop solution (1% 3,3’,5,5’-tetraethylbenzidine, 0.01% H_2_O_2_, 100 mM sodium acetate, 100 mM citric acid). The reaction was stopped after 3 min by adding 0.8 M H_2_SO_4_. Absorbance was measured at 450nm.

### Limited proteolysis

sE2 (0.2mg/ml) was incubated with Endoproteinase GluC (New England Biosciences) at 1:50 (w/w) ratio in TBS at 37°C for up to 4 hours. Proteolysis was halted by boiling in Laemmli buffer at specific intervals and the digestion products identified by SDS-PAGE followed by western blot.

### SDS-PAGE/Western Blotting

Samples were run on MiniPROTEAN 4-12% gels (BioRad, CA, USA) and transferred on to nitrocellulose membrane. The blots were blocked in PBS + 2% milk solution + 0.1% Tween-20 and then probed sequentially with J6.36 mAb and goat anti-mouse secondary conjugated to horseradish peroxidase. Chemiluminescence signal was then measured using a Chemidoc MP (BioRad, CA, USA).

### Circular dichroism spectroscopy

Circular dichroism experiments were performed using the B23 nitrogen-flushed Module B end-station spectrophotometer at B23 Synchrotron Radiation CD BeamLine located at Diamond Light Source. They were performed by the BeamLine lab manager as a mail-in service. Four scans of protein samples in 50 mM NaF and 20 mM NaH_2_PO_4_ and Na_2_HPO_4_ pH 7.2 were acquired in the far-UV region (175 - 260 nm) in 1 nm increments using an integration time of 1 sec, 1 nm bandwidth and pathlength of 0.00146 cm, at 20°C. Results obtained were processed using CDApps v4.0 software (Hussain et al., 2015). The scans were averaged and spectra have been normalized using average amino acid molecular weight which was calculated for each sample. Spectra presented are difference spectra meaning the buffer baseline has been subtracted from the observed spectra, and zeroed between 253 – 258 nm. Secondary structure deconvolution from CD spectra was carried out using the CDApps CONTINLL algorithm and SP29 reference data set referencing 29 soluble protein structures.

### Nano differential scanning fluorimetry

The melting temperature (T _m_) of sE2 (1mg/ml) was calculated using the Prometheus NT.48 (Nanotemper, Munich, Germany) during heating in a linear thermal ramp (1°C, 20 – 90°C) with an excitation power of 30%. The fluorescence at emission wavelengths of 350 and 330 nm was used to determine changes in tyrosine and tryptophan environments. The T_m_ was calculated by fitting the Boltzmann sigmoidal curve to the first derivative of the fluorescence ratios (350 nm/330 nm).

### Mathematical modelling

We applied a mathematical model described in a previous publication (Kalemera et al., 2019) to the novel data generated by this study. Our model uses a series of differential equations to describe the processes via which HCV particles, having bound to the cell membrane, acquire CD81 and SR-B1 receptors. Viruses are modelled as uniformly having N_e_ E2 proteins available to bind cellular receptors. We consider an E2 protein as being either bound or unbound to CD81. If it is unbound to CD81 we consider whether it is bound or unbound to SR-B1; once E2 is bound to CD81 we are unconcerned about its binding to SR-B1. Thus we represent the population as existing on a grid of points M_ij_, indicating viruses that have i copies of E2 bound to CD81 and j copies of E2 bound to SR-B1 but not to CD81. All viruses begin at the point M_00_, then progress through the grid (see Figure 5 in Kalemera et al., 2019).

At the point M_ij_, viruses have N_e_-i-j free E2. We suppose that they bind SR-B1 at some rate s (Figure S4). Viruses bind CD81 via two specific processes. Firstly, viruses bind CD81 via an SR-B1 independent process at the rate c_1_. SR-B1 engagement provides a priming mechanism, which facilitates a second SR-B1-mediated acquisition of CD81, occurring at the rate c_2_. Viruses which gain a pre-specified threshold number of CD81 receptors pass at some rate, e, into the down-stream process of viral entry. Viruses die at constant rate d, which is arbitrarily pre-specified; in our model all other processes occur at some rate relative to this value. Viruses gaining entry to the cell are modelled as being in the state E, while viruses which are dead are modelled as being in the state D.

Our model considers data from a number of states in which the number of CD81 and SR-B1 receptors has been altered (Figure 1C & D). In a given system we model the proportion of CD81 receptors as pc, and the proportion of SR-B1 receptors as ps, where the value in unmodified cells is 1 in each case.

Following the above, we obtain the following equations for viral progress across the membrane:

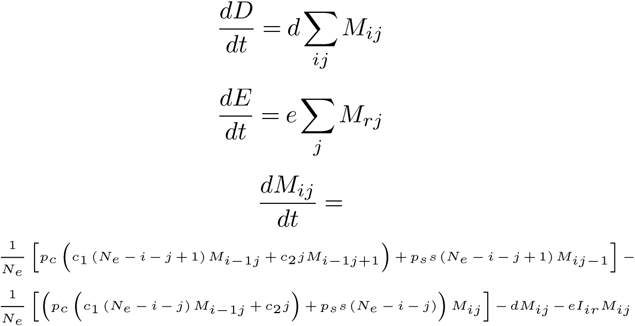

where I_ir_=1 if i=r and I_ir_=0 if ir, and the system has the initial conditions D=0, E=0, M_00_=1, and M_ij_=0 for all i>0 and j>0, and the system is bounded by the constraints 0 i r and 0 j N_e_.

Given a set of input parameters, a fourth-order Runge-Kutte scheme with adaptive step size was used to propagate the system until the sum of terms D+E was greater than 0.999, indicating that 99.9% of the simulated viruses had either died or gained entry. The probability P that a single virus gains entry to a cell was then calculated as

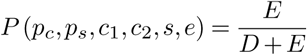

An optimisation procedure was run to fit the model to data from experiments. As in a previous publication, experiments modelling viral entry were performed in replicate. To estimate the values p_c_ and p_s_, indicating the extent of available receptor, we used fluorescence microscopy measurements of receptor blockade/over-expression performed in parallel with the infection experiments, as previously described (Kalemera et al., 2019). These values were normalised to the range 0 to 1 in the case of receptor knockdown experiments, or above 1 in the case of overexpression experiments; 1 being the availability in unmodified cells.

We now consider data describing a level of receptor availability (p_c_, p_s_) and a number of observed foci of infection; using the index i we term the latter value o_i_. Given a knowledge of the number of input particles and the number of cells observed in a well, this can be understood in terms of a probability of a given cell being infected by the virus. Observations were modelled as being distributed according to a double Poisson distribution with mean _i_ and parameter

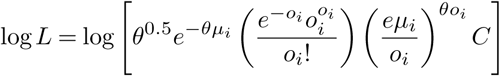

where

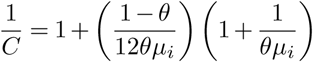

An estimate of the dispersion parameter was calculated by fitting a single value i to each set of values oi arising from the same level of receptor availability with no other constraint on the i. This parameter was then used in the likelihood function to fit values i=niP(pc, ps,c1,c2,s,e), where ni is the number of cells observed in a well and P(pc, ps,c1,c2,s,e) is the probability of viral entry calculated from the differential equation model described above; parameters c1, c2, s, Ne, and e were optimised to give a maximum likelihood fit to the data.

### Molecular dynamic simulations

The complete model of the J6 E2 ectodomain was generated as previously described (Stejskal et al., 2020). For simulations of the mutant glycoprotein, the respective amino acid substitutions were modelled in using Modeller (Webb and Sali, 2016).

We performed MD simulations in explicit solvent using the Amber 16 GPU-based simulation engine (Case et al., 2005). The model was solvated in a truncated octahedral box using OPC water molecules. The minimal distance between the model and the box boundary was set to 12 Å with box volume of 4.2 x 10^5^ Å^3^. Simulations were performed using the ff14SB force field on GPUs using the CUDA version of PMEMD in Amber 16 with periodic boundary conditions. CONECT records were created using the in-house MakeConnects.py script to preserve the disulphide bonds throughout the simulations. MolProbity software was used to generate physiologically relevant protonation states.

Minimisation and equilibration: The systems were minimised by 1000 steps of the steepest descent method followed by 9000 steps of the conjugate gradients method. Sequential 1ns relaxation steps were performed using the Lagevin thermostat to increase the temperature from 0 to 310 K, with initial velocities being sampled from the Boltzmann distribution. During these steps, pressure was kept constant using the Berendsen barostat. All atoms except for the modelled residues, hydrogen atoms and water molecules were restrained by a force of 100 kcal/mol/Å^2^. The restraint force was eventually decreased to 10 kcal/mol/Å^2^ during subsequent 1ns equilibration steps at 310 K.

A further minimisation step was included with 1000 steps of the steepest descent method followed by 9000 steps of the conjugate gradients method with all backbone atoms restrained by a force of 10 kcal/mol/Å^2^. The systems were then subjected to four 1ns long equilibration steps at constant pressure with stepwise 10-fold reduction of restraint force from 10 to 0 kcal/mol/Å^2^. All minimisation and equilibration stages were performed with a 1fs time step.

Production runs: An initial 1*µ*s production run was simulated under constant volume and temperature using the Langevin thermostat. SHAKE was used in all but the minimisation steps; this, in combination with hydrogen mass repartitioning, permitted 4 ft time-steps during the production runs. Short-range cutoff distance for van der Waals interactions was set to 10Å. The long-distance electrostatics were calculated using the Particle Mesh Ewald Method. To avoid the overflow of coordinates, iwrap was set to 1. Default values were used for other modelling parameters. To achieve independent repeat simulations, we performed steps to decorrelate the output from the equilibration process. Initial velocities were generated from the Boltzmann distribution using a random seed. The coordinates, but not velocities, from the final equilibration step were used as input for a short (40ns) production run. The coordinates, but not velocities, from this run were used for a second 4ns production run. This was followed by a 1*µ*s production run. This process was repeated for each independent simulation.

The MD trajectories were analysed using scripts available in cpptraj from Amber Tools 16. For RMSF/RMSD analyses, the average structure generated from the given trajectory was used as the reference structure. The analyses were performed using the backbone C*α*, C and N atoms unless otherwise stated.

### Molecular modeling

Molecular model visualisation was performed with UCSF Chimera (Pettersen et al., 2004).

### Statistical analysis

All statistical analysis was performed in Prism 6.0 (GraphPad, CA, USA). In the majority of cases ordinary one-way ANOVA was performed using Dunnett’s multiple comparison test, using WT virus as a control. Unpaired t-test was performed assuming equal standard deviation using a two-tailed p-value. The F-test was used to compare fitted curves.

## Data availability

All MD trajectories will be made freely available upon final publication.

## ACKNOWLEDGEMENTS

We would like to thank Prof. Greg Towers, Dr. Clare Jolly, Prof. Richard Milne, Dr. Clare Bennett and Prof. Ben Seddon for scientific and academic support, and to colleagues who provided resources (see Methods). We are also grateful to Dr. Colman, Dr. Jackson and Dr. Louis for providing encouragement. This work was supported by Sir Henry Dale Fellowships to J.G. (107653/Z/15/Z) and C.J.R.I. (101239/Z/13/Z); a Wellcome Trust studentship to L.S. (109162/Z/15/Z); a Medical Research Council studentship to M.D.K.. (mrc.ukri.org); C.J.R.I was also supported by Deutsche Forschungsgemeinschaft (DFG, Grant SFB 1310), the Medical Research Council (ref: MC_UU_00002/11) and the Isaac Newton Trust.

## AUTHOR CONTRIBUTIONS

**Conceptualisation**, J.G.; **Methodology**, L.S., M.D.K., W.D.L., D.S.M., T.D., Z.K., C.J.R.I, A.J.S., J.G.; **Investigation**, M.D.K. (Functional experiments with HCVcc/pp/sE2), L.S. (Computational biology and biophysical experiments), M.P., L.W., M.K-V., C.J.R.I, J.G.; **Resources**, W.R.; **Writing**, J.G., M.D.K.; **Funding Acquisition**, L.S., M.D.K., C.J.R.I, J.G.; **Supervision**, A.J.S., J.G..

