## Supplementary Information for "An Entropic Safety Catch Controls Hepatitis C Virus Entry and Antibody Resistance"

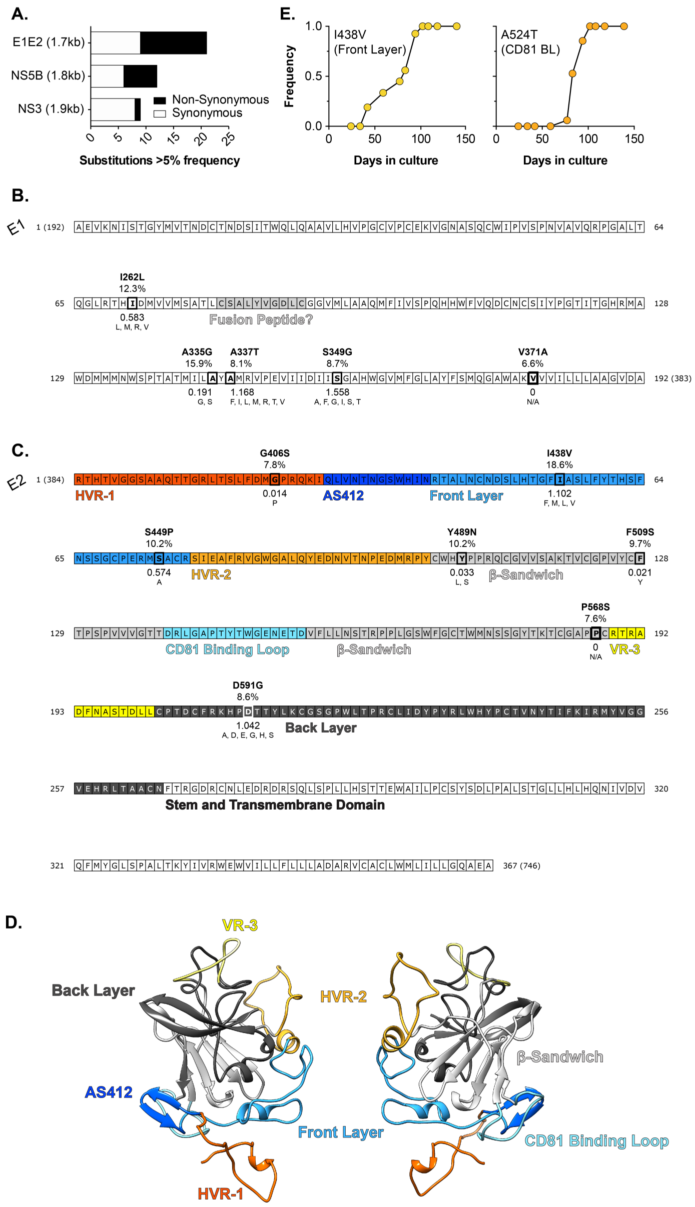


**Figure S1 Location and frequency of non-synonymous substitutions in E1E2.** J6/JFH HCVcc was continuously propagated in Huh-7.5 cells for 20 weeks with viral evolution monitored by NGS. **A.** The proportion of synonymous (white) and non-synonymous (black) substitutions occurring, at day 42, at >5% frequency in three coding regions of comparable length: E1E2 (entry glycoproteins), NS5B (polymerase), NS3 (protease). **B. & C.** Non synonymous substitutions mapped onto linear representations of the E1 (B.) and E2 (C.) protein sequences. Functional and structural features are highlighted (e.g. the putative fusion peptide). The E2 sequence is color-coded by region, as denoted on the three-dimensional model (**D.**). Each substitution is annotated with: i) the observed mutation, ii) the frequency in the experimental population at day 42 (%), iii) the shannon entropy calculated from patient derived sequences, with low values denoting high conservation iv) substitutions that naturally occur at these positions in patient derived sequences; N/A (not applicable) indicates the lack of natural variants. **E.** Emergence and fixation of substitutions at position 438 (Front layer) and 524 (CD81 Binding Loop); by 100 days all viruses possessed the I438V A524T double mutation. Note that residue numbering is relative to the start of the HCV polyprotein and corresponds to the position in the H77 reference strain, as is convention within the HCV field.

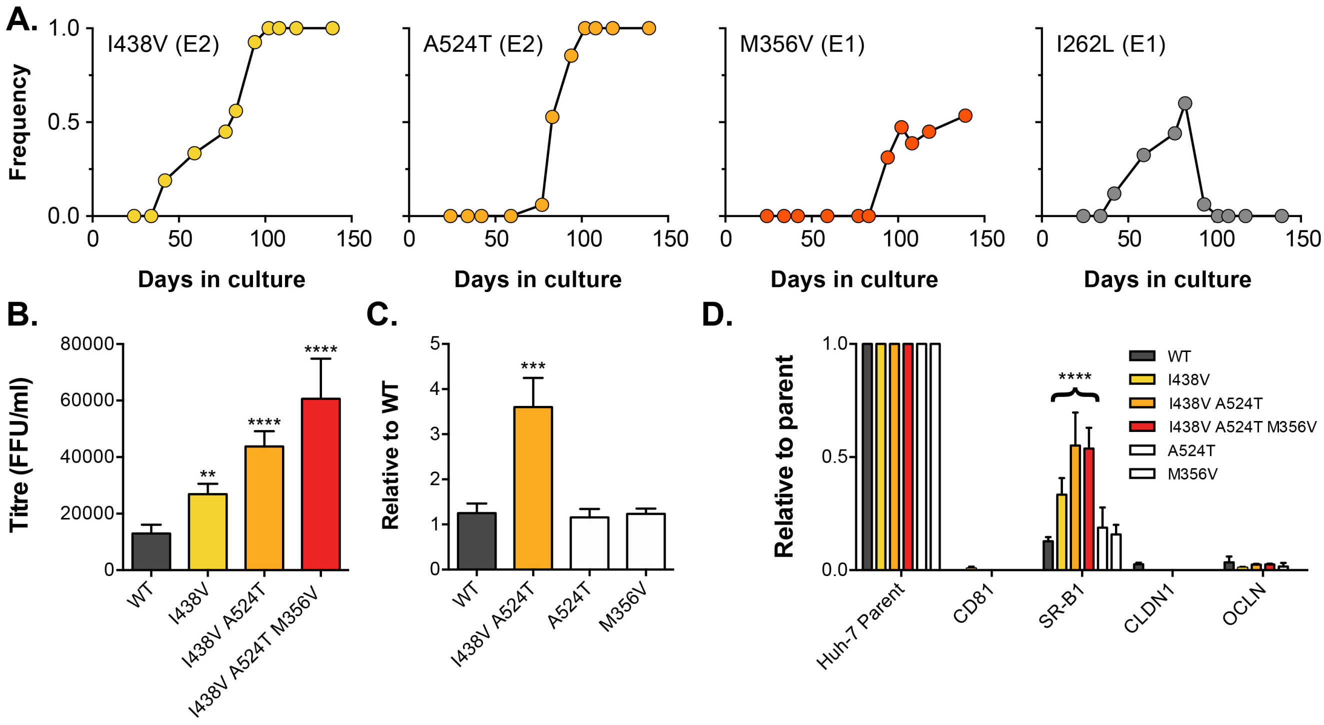


**Figure S2 The emergence of a mutant lineage with altered receptor dependency.** J6/JFH HCVcc was continuously propagated in Huh-7.5 cells for 20 weeks with viral evolution monitored by NGS. **A.** The frequency of example mutations throughout the culture experiment. By day 100, all viruses possessed the I438V A524T E2 mutations, ~50% had an additional mutation in E1 (M356V), whilst a previously prevalent mutation (I262L) had been lost from the population. **B.** Infectivity of WT; I438V; I438V A524T and I438V A524T M356V HCV, expressed as foci forming units per ml. Values were normalised for input particle numbers. **C.** Infection of WT, I438V A524T and stated single mutants. Data is expressed relative to WT infection. **D.** WT and mutant HCV infection of parental Huh-7 cells and those CRISPR/Cas9 engineered to knock out the stated HCV entry factor. To aid direct comparison of each mutant, infection values have been expressed relative to that observed in parental Huh-7 cells. Error bars indicate standard error of the mean, asterisks denote statistical significance (ANOVA, GraphPad Prism).

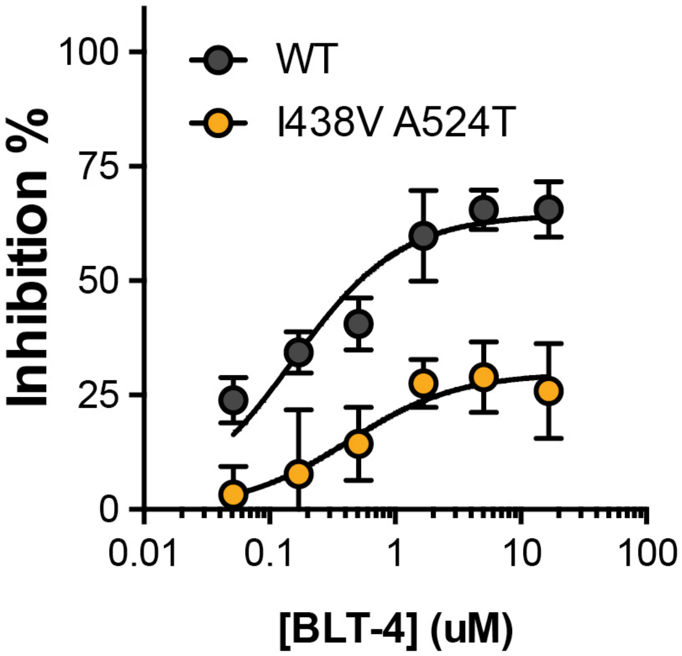


**Figure S3 Inhibition of HCVcc infection by BLT-4.**
Huh-7.5 cells, treated with a serial dilution of a small molecule inhibitor of SR-B1 (BLT-4), were infected with WT and mutant HCVcc. Infection is expressed as % inhibition relative to untreated cells. Data points represent the mean of three independent experiments. The data was fitted with a hyperbola function and the curves were determined to be statistically significant (F-test, GraphPad Prism), error bars indicate standard error of the mean.

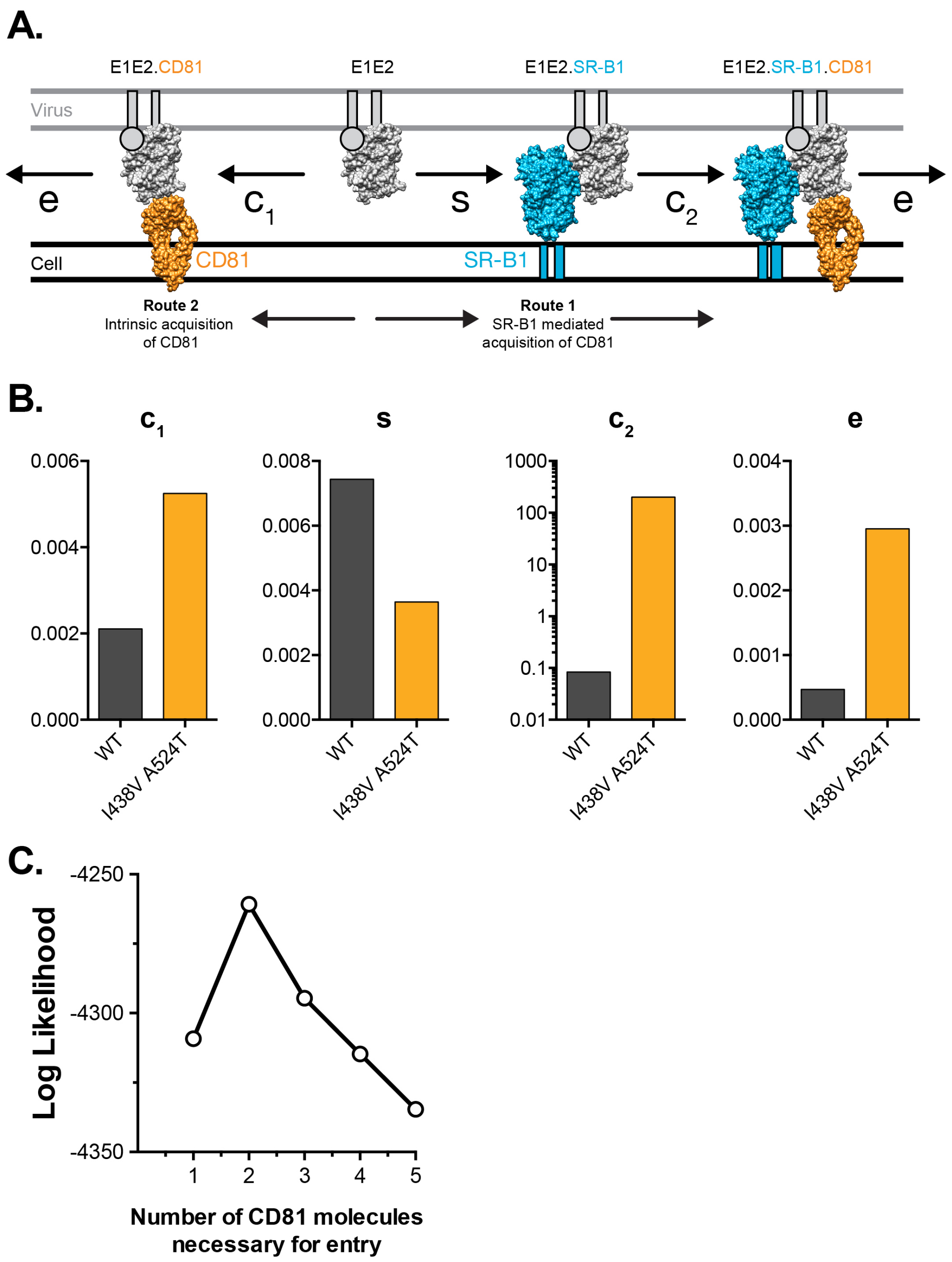


**Figure S4 Mathematical modelling of HCV entry.**

We previously developed a mathematical model to explore HCV receptor engagement and entry (Kalermera et. al. 2019). **A.** E1E2-receptor interactions at the cell surface, as recreated in our model. To achieve entry E1E2 must acquire CD81, this can occur via two routes. Route 1 SR-B1 mediated: prior binding to SR-B1 (at rate ***s***), primes E1E2 for interaction with CD81 (at rate $\boldsymbol{c_2}$). Route 2 intrinsic binding: E1E2 interact with CD81 without prior engagement of SR-B1 (at rate $\boldsymbol{c_1}$). Once sufficient molecules of CD81 have been acquired the virus particle proceeds along the entry pathway (including endocytosis and fusion) at rate ***e***. Molecular cartoons are based on previously published structures and are drawn to scale. **B.** Receptor dependency data from WT and mutant HCVcc (Figure 2) was integrated into the mathematical model, as previously described, allowing estimation of the parameters defined above. Plots display WT (grey) and I438V A524T (orange) values for each rate constant. **C.** The model was used to explore the stoichiometry of CD81 engagement during HCV entry; the plot displays likelihood values for 1-5 molecules of CD81. Peak likelihood occurs at 2 molecules.

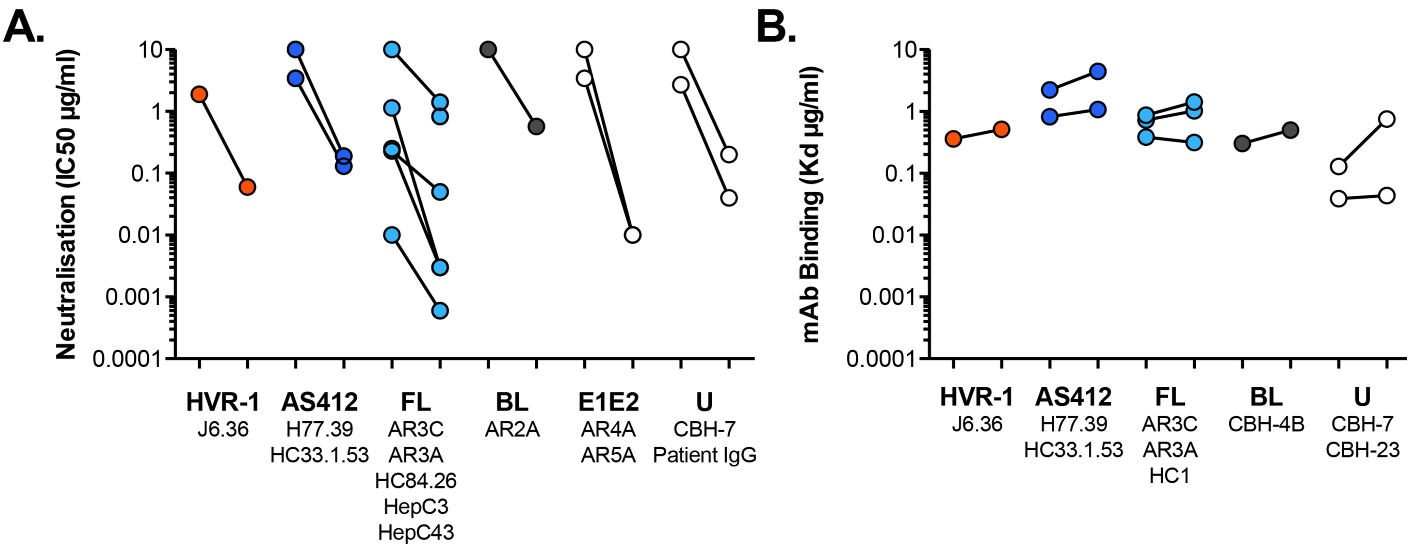


**Figure S5 Antibody sensitivity and antigenicity of WT and I438V A524T HCV.**
**A.** 50% inhibitory concentrations of mAbs (as provided in Figure 3D) separated according to antigenic targets. mAb names are provided for each target. FL and BL refer to Front Layer and Back Layer. E1E2 mAbs bind to a discontinuous epitope comprising elements of both E1 and E2. U represents poorly defined epitopes. **B.** Estimated dissociation constants (as provided in Figure 3F), separated according to antigenic targets.

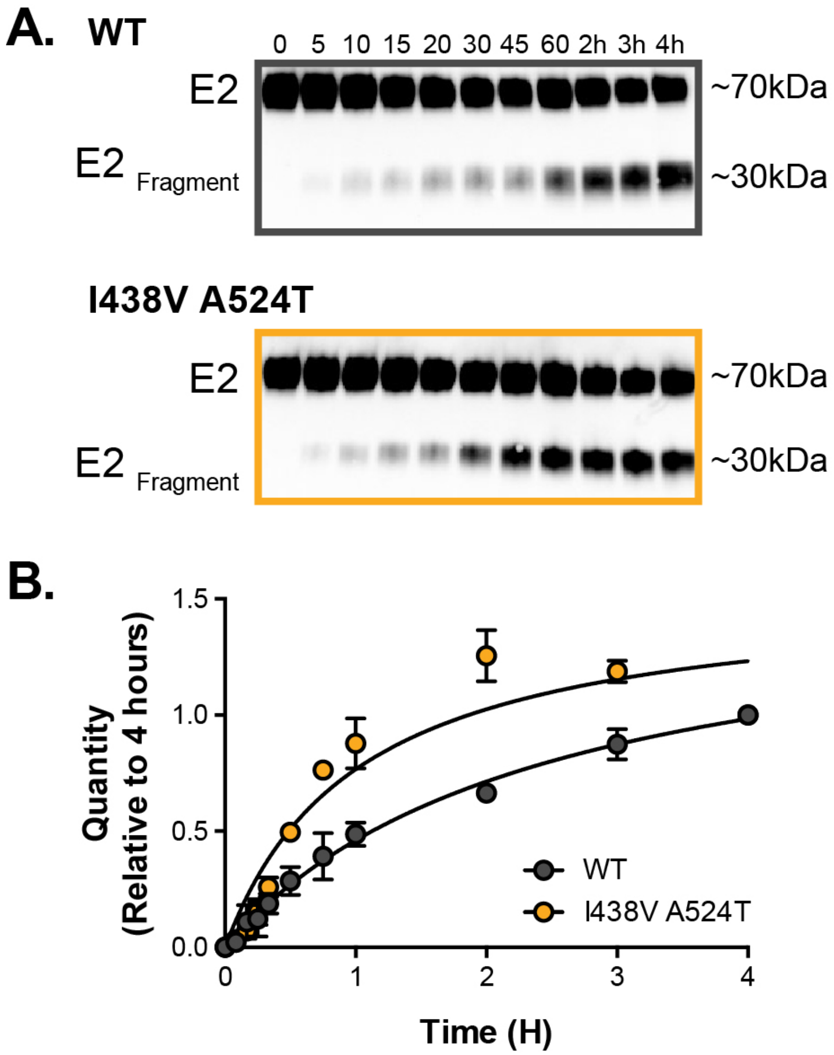


**Figure S6 Limited proteolysis of soluble E2.**
Soluble E2 (WT or I438V A524T) was incubated at 37°C with endproteinase GluC for up to 4 hours. **A.** Representative western blot images of digestion over time (minutes and hours as indicated); incubation with GluC gives rise to a ~30kda digest product ($E2_{Fragment}$). **B.** The kinetics of digestion was evaluation by measuring the quantity of $E2_{Fragment}$; the data is expressed relative to the intensity of product at 4 hours, data points represent the mean of three independent experiments, error bars indicate standard error of the mean. Data were fitted with a hyperbola function and curves were compared to determine statistical significance (F-test, GraphPad Prism).

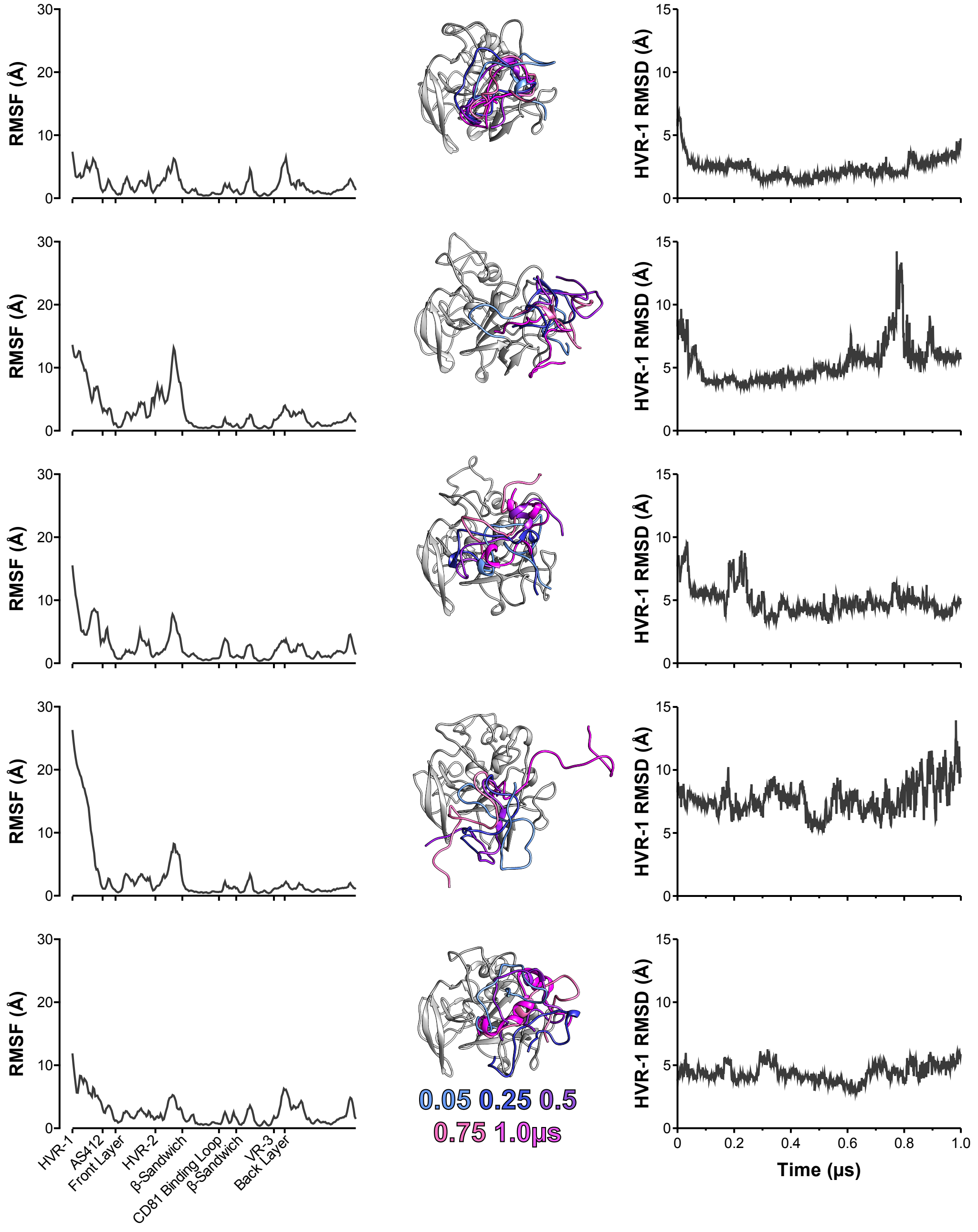


**Figure S7 Five independent 1μs MD simulations of WT E2.**

The conformational dynamics of WT E2 ectodomain were explored by MD simulations. E2 RMSF (left), images illustrating HVR-1 mobility (middle), and HVR-1 RMSD (right) are provided for each simulation.

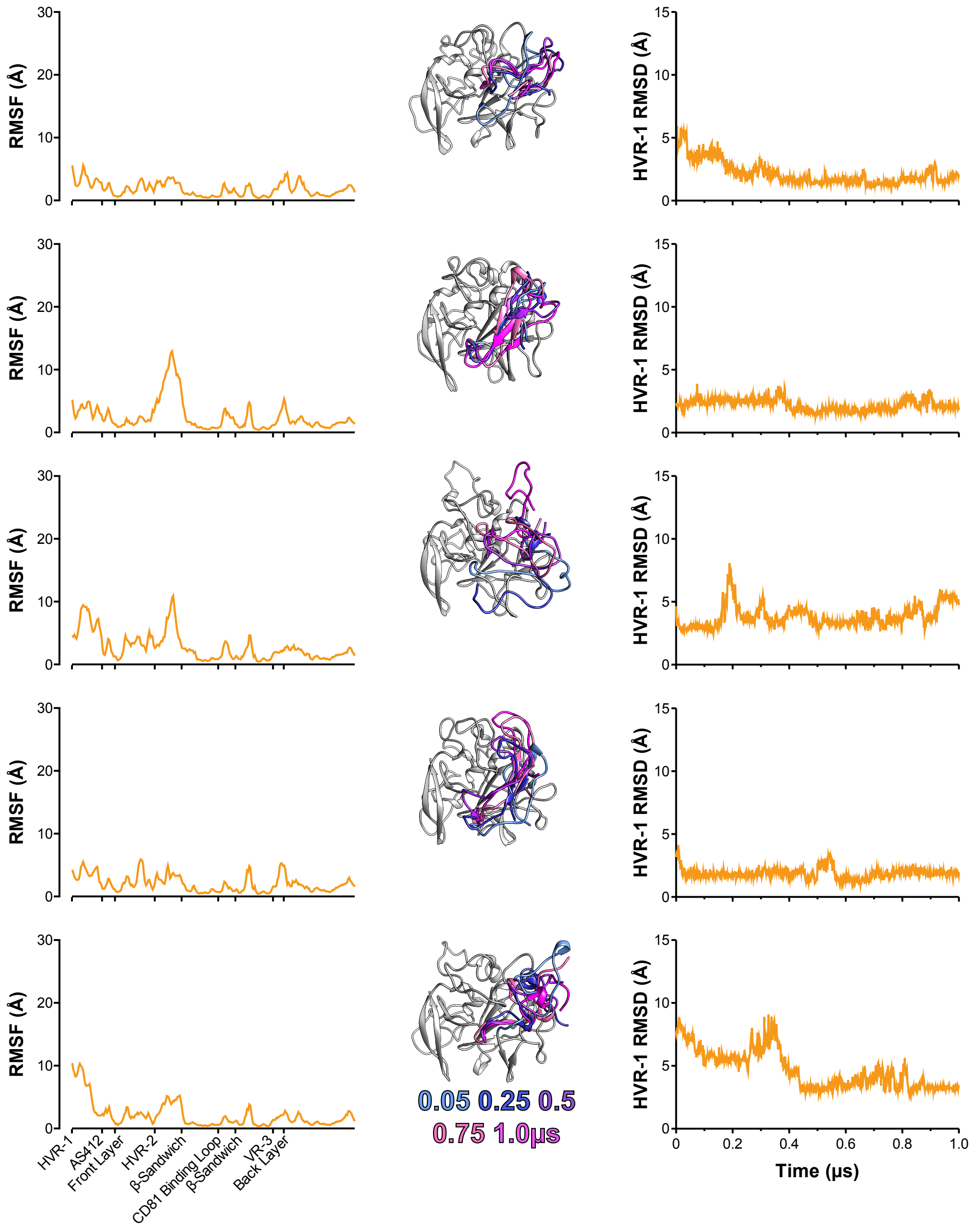


**Figure S8 Five independent 1μs MD simulations of I438V A524T E2.**
The conformational dynamics of I438V A524T E2 ectodomain were explored by MD. E2 RMSF (left), images illustrating HVR-1 mobility (middle), and HVR-1 RMSD (right) are provided for each simulation.

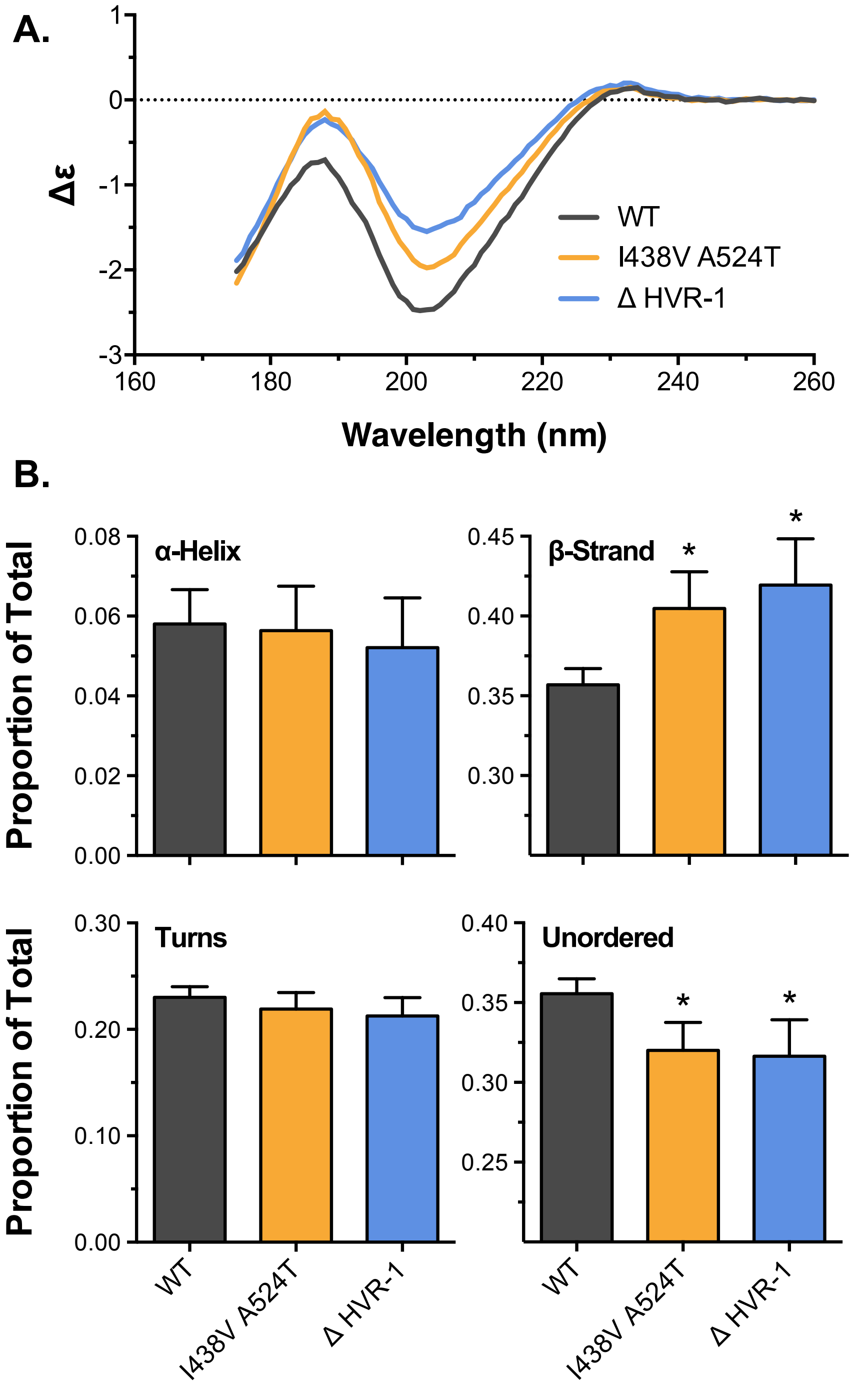


**Figure S9 Biophysical analysis of E2 by circular dichroism spectroscopy.**
**A.** Circular dichroism spectra for WT, I438V A524T and ΔHVR-1 soluble E2. **B.** Estimates of the secondary structure content found in E2, data is expressed as a proportion of total. Asterisks indicate statistical significance (ANOVA, Graphpad, Prism).

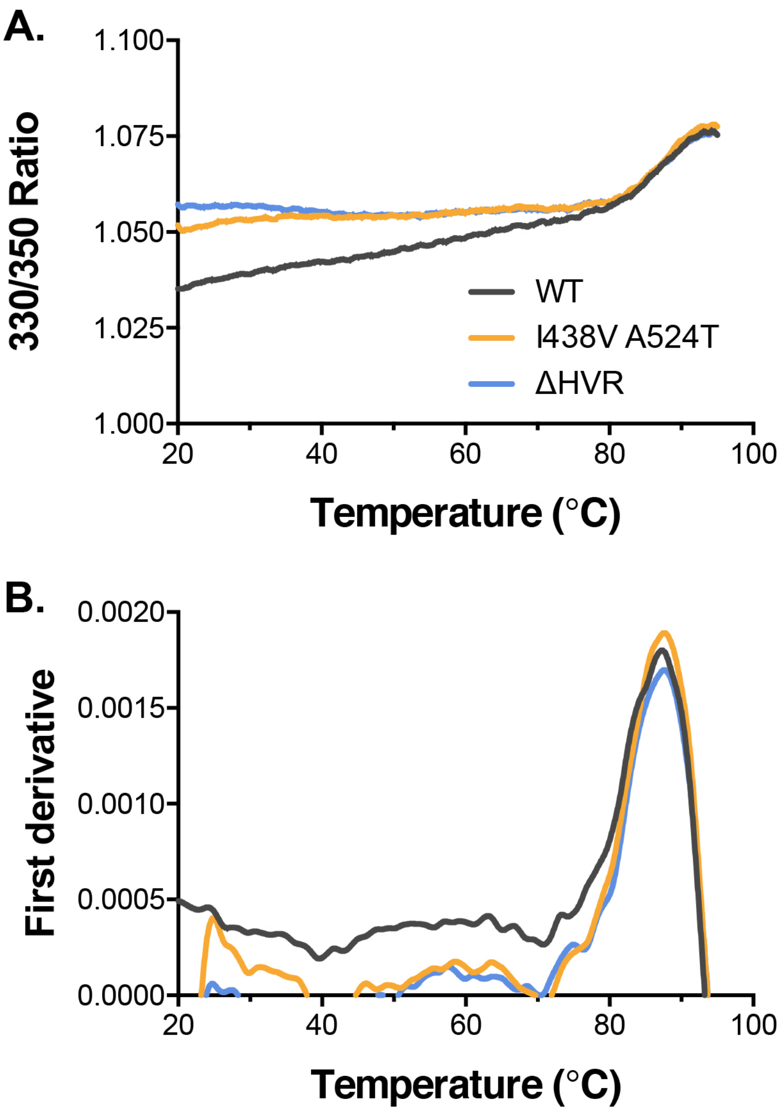


**Figure S10 Biophysical analysis of E2 by nanoscale differential scanning fluorimetry.** **A.** Intrinsic fluorescence ratio (330nm over 350nm) for WT, I438V A524T and ΔHVR-1 soluble E2 at increasing temperatures. **B.** First derivative transformation of fluorescence ratio; peak indicates protein denaturation.

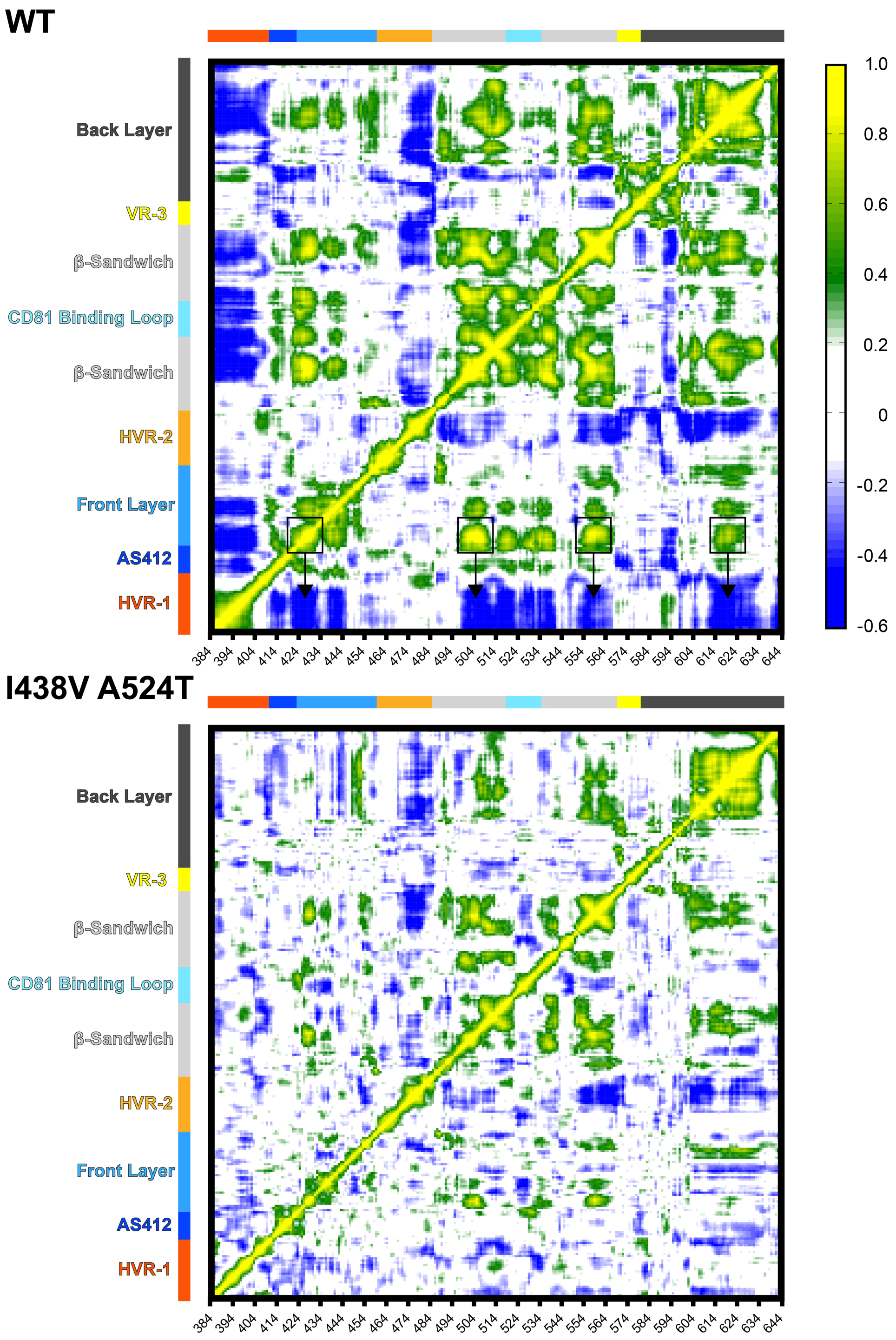


**Figure S11 Dynamic cross-correlation analysis of E2 MD simulations.**
Dynamic cross correlation provides a residue-by-residue pairwise comparison of motion in MD trajectories to reveal correlations/anti-correlations in protein movement. Average DCC matrices for WT and I438V A524T E2 MD simulations are provided. As depicted in the key, yellow/green indicate positive correlations; blue indicates negative correlations; white indicates lack of correlation. The black boxes indicate hotspots of correlation at disulphide bonds; these correspond to areas in which HVR-1 motions are felt most strongly (arrows).

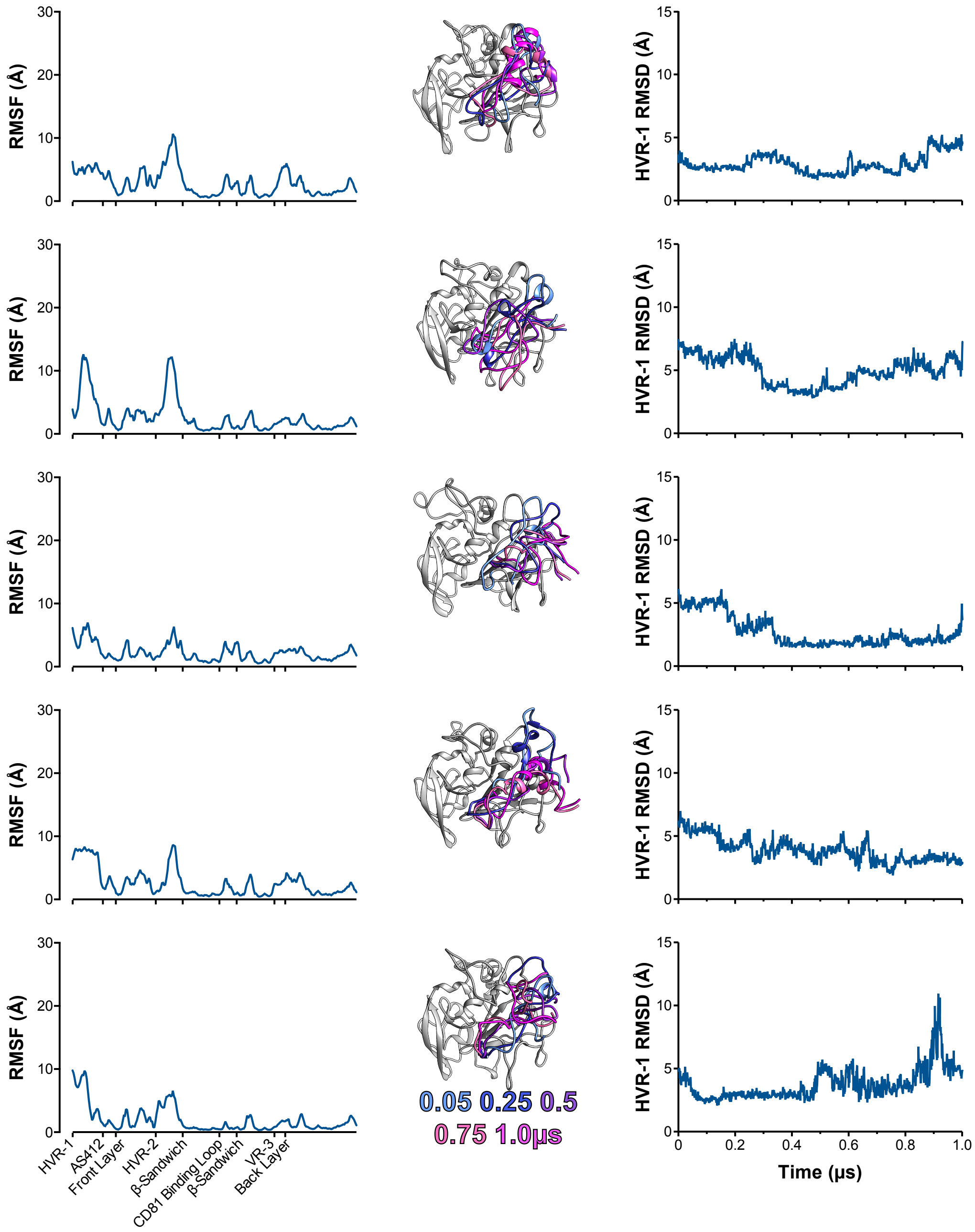


**Figure S12 Five independent MD simulations of S449P E2.**

The conformational dynamics of S449P E2 ectodomain were explored by MD. E2 RMSF (left), images illustrating HVR-1 mobility (middle), and HVR-1 RMSD (right) are provided for each simulation.

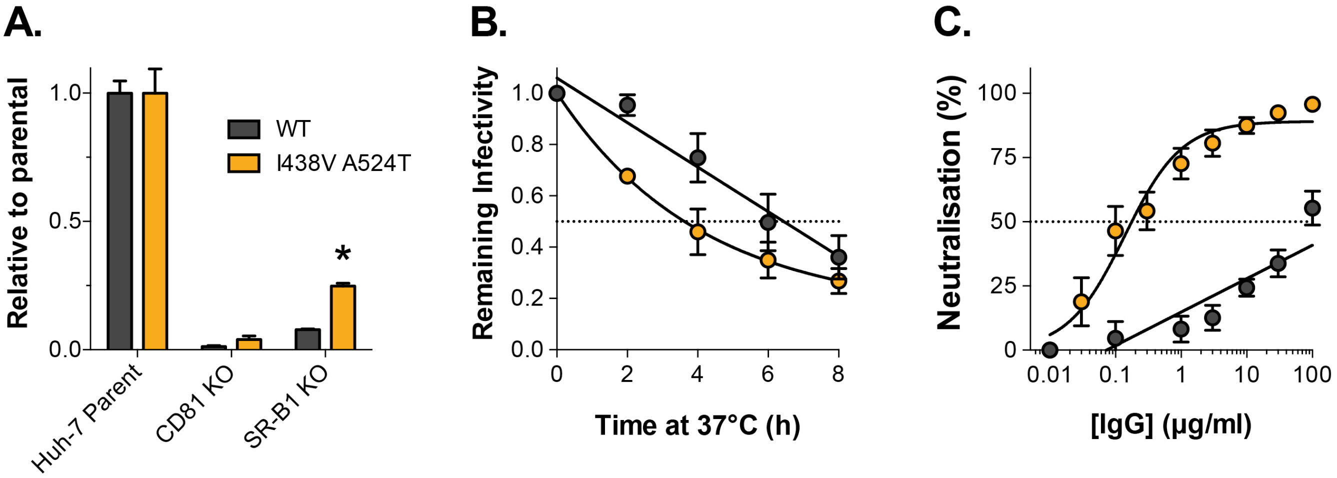


**Figure S13 Hyper-reactive phenotype is recapitulated in HCV pseudoparticles**

We characterised WT and I438V A524T E1E2 in the context of HCV pseudoparticles. **A.** HCVpp infection of parental or receptor knockout Huh-7, data is expressed relative to parental cells. **B.** Stability of HCVpp at 37°C, data points represent the mean of three independent experiments, data was fitted using an exponential decay function. **C.** Neutralisation of HCVpp by patient IgG, data points represent the mean of three independent experiments. Data was fitted with a hyperbola function (I438V A524T) or semilog function (WT). In each plot error bars indicate standard error of the mean; asterisks denote statistical significance (ANOVA, Graphpad prism); all curves determined to be statistically significant (F-test, Graphpad prism).
